# Phosphorylation by CK2 increases the SUMO-dependent activity of Cytomegalovirus transactivator IE2

**DOI:** 10.1101/655282

**Authors:** Vasvi Tripathi, Kiran Sankar Chatterjee, Ranabir Das

## Abstract

Viral factors manipulate the host post-translational modification (PTM) machinery for replication. Distinctly, phosphorylation and SUMOylation can regulate the activity of human cytomegalovirus (HCMV) protein IE2. However, the molecular mechanism of this process is unknown. Taking a structural, biochemical and cellular approach, we uncover a cross-talk of phosphorylation and SUMOylation exploited by IE2. A scan for the SUMO Interacting Motifs (SIMs) revealed two SIMs in IE2. A real-time SUMOylation assay indicated that the N-terminal SIM (IE2-SIM1) enhanced IE2 SUMOylation up to 4-fold. Kinetic analysis and structural studies proved that IE2 is a SUMO cis-E3 ligase. Two putative CK2 sites adjacent to IE2-SIM1 are phosphorylated *in-vitro* and in cellular conditions. Phosphorylation drastically increased the IE2/SUMO affinity, IE2-SUMOylation and cis-E3 activity of IE2. Additional salt-bridges between the phosphoserines and SUMO account for the higher IE2/SUMO affinity. Phosphorylation also enhances the SUMO-dependent transactivation activity and auto-repression activity of IE2. Together, our findings highlight a novel mechanism where SUMOylation and phosphorylation of the viral cis-E3 ligase and transactivator protein IE2, works in tandem to enable transcriptional regulation of viral genes.

**Author summary:** The host protein SUMO is a crucial regulator of cellular processes. Conjugation of other proteins to SUMO by a process called SUMOylation, can change the protein’s function or localization and regulate downstream cellular events. The SUMO pathway is exploited by viruses to transcribe viral genes and replicate the viral genome. IE2 is an essential gene of human Cytomegalovirus (HCMV), which acts as a transactivator and helps to transcribe other viral proteins required for viral genome replication and viral assembly. SUMOylation of IE2 is necessary for its function. Here, we have uncovered that IE2 functions as a cis-SUMO-E3 ligase, where a SUMO-Interacting Motif (SIM) in IE2 enhances its SUMOylation. Interestingly, phosphorylation of the SIM in IE2 augments its cis-E3 activity to further increase SUMOylation. Moreover, SIM phosphorylation also enhances the interaction between IE2 and SUMOylated binding partners. Thus, we uncover an exciting process, where phosphorylation enhances both covalent and non-covalent interaction of a protein (IE2) and SUMO. We also observe that the cross-talk of phosphorylation and SUMOylation has significant effects on the transactivation function of IE2. Overall, we discover how a viral protein IE2 exploits crosstalk between SUMOylation and Phosphorylation to enhance its activity and in turn, ensure efficient viral replication.

## Introduction

The Immediate Early (IE) genes are the first viral genes transcribed during infection. The IE gene products are essential to optimize the host cell environment for replication of the viral genome and the transcription of early and late genes. IE1 and IE2 are the two predominant IE proteins in Human Cytomegalovirus (HCMV), which are splicing products of the Major Immediate Early Promoter (MIEP) (Stenberg, Thomsen and Stinski, 1984). While IE1 is essential only at low MOI, IE2 is strictly indispensable for viral replication and growth (Marchini, Liu and Zhu, 2001).

IE2 regulates HCMV growth at multiple levels. It arrests the cell cycle in ‘pseudo-S’ phase at the G1-S boundary to facilitate viral replication over host genome replication (Petrik, Schmitt and Stinski, 2006). IE2 is an essential factor for replication complex assembly (Colletti *et al*., 2004). Finally, IE2 works as transactivator for early/late viral genes and various host genes (Hiagemeier *et al*., 1992). IE2 transactivates TATA Box containing promoters with the help of basal transcription factors (TBP, TFIIB, and TAFs) (Hiagemeier *et al*., 1992; Bryant *et al*., 2000). Interestingly, IE2 regulates its promoter MIEP by binding to cis-regulatory sequence downstream of the transcription start site (Hoffmann *et al*., 1993). Apart from transcription factors, IE2 also functions with transcription coactivators and chromatin modifiers to regulate the host transcriptional machinery (Lang *et al*., 1995; Bryant *et al*., 2000; Reeves *et al*., 2006; Lee *et al*., 2011).

IE2 is a pleiotropic regulator. Hence, its expression and activity is tightly regulated transcriptionally, post-transcriptionally, and also by Post Translational Modifications (PTMs) like SUMOylation and phosphorylation (Hoffmann *et al*., 1993; Ahn *et al*., 2001; Heider *et al*., 2002; Arend, Lenarcic and Moorman, 2018). SUMOylation is the covalent addition of protein Small Ubiquitin-like Modifier (SUMO) to the lysine side chain of a substrate (Gareau and Lima, 2010). SUMOylation occurs through an enzymatic cascade involving sequential action of an E1 activating enzyme, an E2 conjugating enzyme (UBC9), and a few E3 ligases. SUMOylation of IE2 occurs at the two lysine residues K175 and K180 (Hofmann, Flo and Stamminger, 2000; Ahn *et al*., 2001). The Cytomegalovirus with SUMOylation-deficient IE2 has severe growth defects due to impaired initiation of gene expression, replication compartment assembly and MIEP autoregulation (Berndt *et al*., 2009; Reuter *et al*., 2018). Proteins with a SUMO Interacting Motif (SIM) can identify SUMO or SUMOylated proteins via the noncovalent SIM/SUMO interaction (Song *et al*., 2004). IE2 also contains a SIM at its N-terminus, which is essential for its localization to nuclear puncta and transactivation activity (Kim *et al*., 2010). The SIM recruits SUMOylated transcription factors (e.g., TAF12) to facilitate transactivation. Moreover, deletion of the SIM decreases IE2 SUMOylation (Berndt *et al*., 2009; Kim *et al*., 2010). Despite the functional importance of IE2/SUMO noncovalent interaction, its molecular details are unknown. Additionally, the molecular mechanism underlying the role of SIM in SUMOylation of IE2 is unclear.

IE2 is phosphorylated by several host kinases (Barrasa, Harel and Alwine, 2005). Phosphorylation by ERK2 (a MAP kinase) inhibits its transactivation activity without affecting auto-repression of MIEP (Harel and Alwine, 1998; Heider *et al*., 2002). IE2 is also phosphorylated by the kinase Casein Kinase 2 (CK2), which is intriguing because CK2 is a part of the viral tegument (Varnum *et al*., 2004). During infection, the uncoating of HCMV tegument releases the tegument CK2 in the host cell to activate MIEP expression (Nogalski *et al*., 2007). CK2 phosphorylates IE2 at the serine-rich region (aa: 258-275) and in the so-called fragment 5B region (aa:180-252, Figure 1A) (Barrasa, Harel and Alwine, 2005). Phosphorylation of the serine-rich region depletes the transactivation activity of IE2 (Barrasa, Harel and Alwine, 2005). However, the sites of phosphorylation within fragment 5B and the effect of this phosphorylation on the activity of IE2 is unknown.

**Figure 1.**
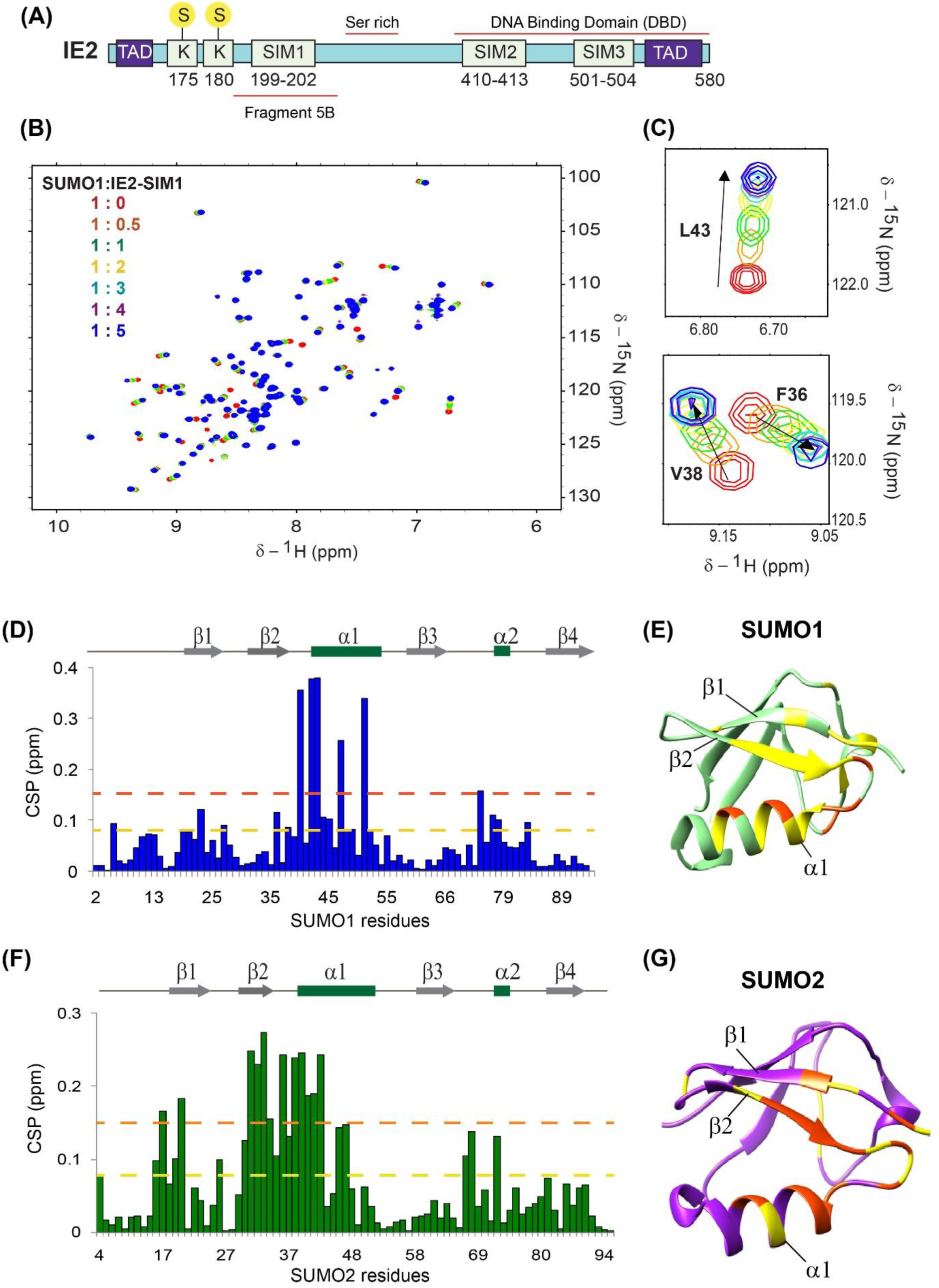
Interactions between IE2-SIM1 and SUMO. (A) A schematic of the domains in IE2. The three predicted SIMs, and the two SUMOylation site lysines K175 and K180 are shown. Yellow circles marked ‘S’ denote SUMO. The transactivation domains are shown as blue boxes and marked as ‘TAD.’ The Fragment 5B, Serine rich region, and DBA binding domain are shown. (B) Overlay of the ^15^N-edited HSQC spectra of free ^15^N-SUMO1 (red) with different stoichiometric ratios of IE2-SIM1 as given in the top left-hand side of the spectra. (C) Two regions of the spectra are expanded to show a shift of SUMO1 resonances during titration. (D) The CSPs in SUMO1 upon binding to IE2-SIM1. The chemical shift perturbations (CSP) between the free and the bound form are calculated as CSP = [(δ^H^_free_ – δ^H^_bound_)^2^+ ((δ^N^_free_ – δ^N^_bound_)/5)^2^]^1/2^, where δ^H^ and δ^N^ are the chemical shift of the amide hydrogen and nitrogen, respectively. The yellow and red dashed lines indicate 1 x standard deviation and 2 x standard deviation, respectively. The secondary structure alignment of SUMO1 against its sequence is provided above the plot. The residues with CSPs significantly above the dashed lines are present at the interface of the SUMO1/IE2-SIM1 complex. (E) The significant CSPs are mapped onto the SUMO1 structure. The residues with CSP above yellow and red lines are colored in yellow and red, respectively. (F) The CSPs in SUMO2 upon binding to IE2-SIM1. (G) The significant CSPs mapped on the SUMO2 structure. The residues with CSP above yellow and orange lines are colored in yellow and orange, respectively.

We carried out a study of the predicted SIMs in IE2 to uncover a new SIM at the C-terminus of IE2 by NMR. NMR also confirmed the previously known N-terminal SIM (IE2- SIM1). The titrations indicate that IE2-SIMs bind to both SUMO1and SUMO2 with similar affinity. Adjacent to the N-terminal SIM (IE2-SIM1) are the two SUMOylation sites K175 and K180. We developed a fluorescence-based real-time SUMOylation assay, which confirmed that the IE2-SIM1 enhances SUMOylation of IE2. However, the assay also indicated paralog specificity in SUMOylation of IE2. Kinetic analysis revealed that IE2 functions as a SUMO cis-E3, where the IE2-SIM1 reduces K_m_ between UBC9∼SUMO and the IE2 SUMOylation sites to increase the rate of IE2 SUMOylation.

We further carried out a comprehensive study of IE2 phosphorylation in cellular conditions. We report that two putative CK2 phosphorylation sites Ser203 and Ser205 in fragment 5B are indeed phosphorylated in cells, as well as *in-vitro* by CK2. Ser203 and Ser205 are adjacent to the IE2-SIM1. Phosphorylation of Ser203 and Ser205 increases the SUMO/IE2-SIM1 affinity by 8-fold. Structures of the complex between SUMO1/2 and phosphorylated IE2-SIM1 attribute the increased affinity to additional salt-bridges formed between the positively charged residues in SUMO1/2 and the negatively charged phosphate moiety in the phosphoserines. The drastic increase in SIM/SUMO affinity also enhances the SUMO cis-E3 activity, transactivation activity and auto-repression activity in IE2. Together, the HCMV protein IE2 uniquely exploits a cross-talk of two post-translational modifications; phosphorylation and SUMOylation, to regulate transcription of the viral genes and ensure a productive infection.

## Results

### The N-terminal and C-terminal SIMs in IE2 interact with SUMO1 and SUMO2

Bioinformatics analysis using JASSA (Beauclair and Bridier-nahmias, 2015) predicted three SIMs in IE2, SIM1: 199-202, SIM2: 410-413 and SIM3: 501-504 as shown in Figure 1A. Among these, SIM1 was previously identified as a bonafide SIM by pull-down assays in cells (Berndt *et al*., 2009). However, the rest of the SIMs were not investigated before. The affinity of the SIM/SUMO interaction is typically weak, which is difficult to capture by pull-down experiments. Alternately, NMR spectroscopy can detect interaction over a broad range of affinities including weak interactions (Vaynberg and Qin, 2006). We designed peptides corresponding to the putative SIMs and tested the binding of the SIMs to SUMO1 by NMR (Table 1). Perturbations due to the altered chemical environment upon ligand binding are reflected in a shift of the backbone amide resonances in the ^15^N-edited HSQC spectra. The IE2-SIM1 peptide was titrated into a sample of ^15^N-isotope labeled SUMO1, and a series of ^15^N-edited HSQC experiments monitored its effect on ^15^N-SUMO1. An overlay of the HSQCs is given in Figure 1B, and two expanded areas of the HSQC are plotted in figure 1C. A subset of SUMO1 peaks shifted consistently with increasing concentrations of IE2-SIM1. The chemical shift perturbations (CSP) against SUMO1 residues given in Figure 1D, indicated that the maximum perturbations occurred in the residues 35 to 55, which includes the β-strand β2, α-helix α1 and the loop between them (Figure 1E). The interface corresponds to the canonical interface observed in other SUMO/SIM complexes (Song *et al*., 2004). The same titration experiments were repeated for IE2-SIM2 and IE2-SIM3 domains in separate experiments. IE2-SIM3 but not IE2-SIM2, bound to SUMO1 (Figure S1A). The pattern of CSP in SUMO1 upon binding IE2-SIM3 is similar to IE2-SIM1, indicating that both the SIMs bind at the same interface.

**Table 1:**
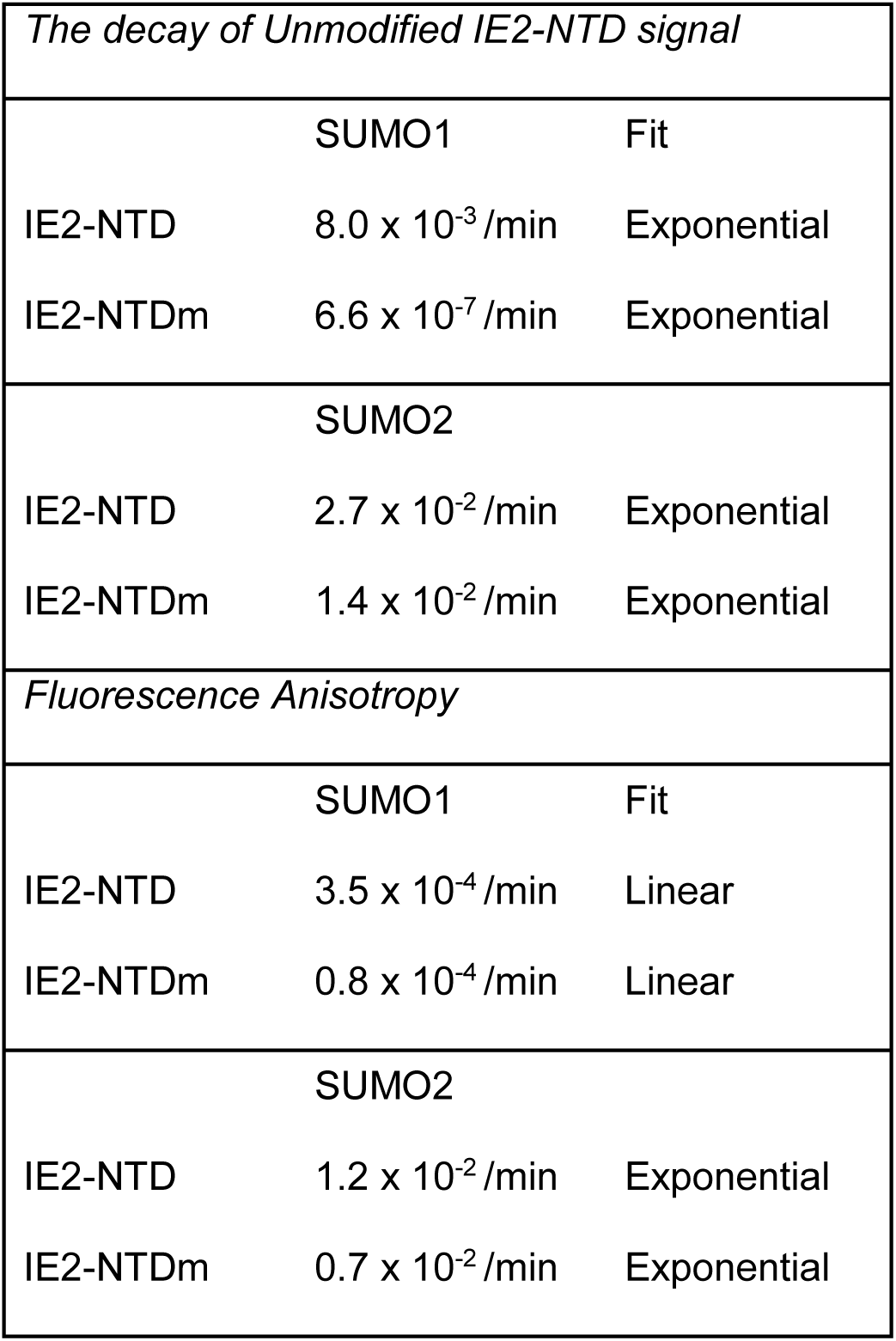
Analysis of possible SUMO Interaction motifs in IE1 and IE2

We repeated the titration experiments with IE2-SIMs against ^15^N-labeled SUMO2 and monitored the effects using ^15^N-edited HSQC spectra of ^15^N-SUMO2. Similar to SUMO1, a significant subset of SUMO2 peaks shifted upon titration with IE2-SIM1 as shown in Figure S1B and S1C. The most perturbed region is between β2 and α1 of SUMO2, which is the known SUMO2/SIM interface (Figure 1F, 1G). Additionally, IE2-SIM3 but not IE2-SIM2, interacted with SUMO2 (Figure S1D).

The CSPs observed during titration were fit against peptide: protein concentration to yield the dissociation constant (K_d_) of the complex (Figure S2, S3, Table 1). The K_d_ of N-terminal IE2-SIM1 was 6-fold lower than the C-terminal IE2-SIM3, indicating that the N-terminal SIM has a higher affinity for SUMO than the C-terminal SIM. Both IE2-SIM1 and IE2-SIM3 do not have paralog specificity and have similar affinities for SUMO1 and SUMO2. Taken together, while IE2-SIM1 and IE2-SIM3 interacted with both SUMO1 and SUMO2, the interaction between IE2-SIM1 and SUMO1/2 was stronger. Hence, the IE2-SIM1 was studied further.

### Casein Kinase II phosphorylates two serines adjacent to IE2-SIM1

IE2 phosphorylation modulates its transactivation activity (Barrasa, Harel and Alwine, 2005). Previous phosphorylation studies have focused on specific sites or domains in IE2 (Harel and Alwine, 1998; Heider *et al*., 2002; Barrasa, Harel and Alwine, 2005). A comprehensive report of phosphorylation sites in the cellular condition is missing. Thus, we studied the sites of IE2 phosphorylation in HEK293T cells taking a proteomics approach, (Figure 2A, Figure S4). Interestingly, in addition to various newly identified phosphorylation sites, two serines adjacent to IE2-SIM1, Ser203, and Ser205, were phosphorylated in cells (Figure 2A). Ser203 and Ser205 are putative Casein Kinase II (CK2) target phosphorylation sites. An *in-vitro* phosphorylation assay of IE2-SIM1 by CK2 was carried out to confirm if these sites are indeed modified by CK2 (Figure 2B). IE2-SIM1 peptide does not include any serines apart from Ser203 and Ser205. It consists of a threonine, which is not the putative CK2 phosphorylation site. Phosphorylation of IE1-SIM1 was carried out using γ-ATP, resolved on SDS-PAGE and detected by autoradiography. As shown in Figure 2C, phosphorylated IE2- SIM1 was observed in the reaction with active CK2, while the heat inactivated CK2 could not phosphorylate IE1-SIM1. IE2-ppSIM1 was further analyzed by MALDI-TOF to find out if it was phosphorylated at a single serine or both the serines (Figure 2D). While mass (m/z) for IE2-SIM1 was 2098 Da, the same for phosphorylated IE2-SIM1 was 2258 Da. The difference of 160Da corresponds to the addition of two phosphate groups, indicating that indeed CK2 phosphorylates IE2-SIM1 at Ser203 and Ser205.

**Figure 2.**
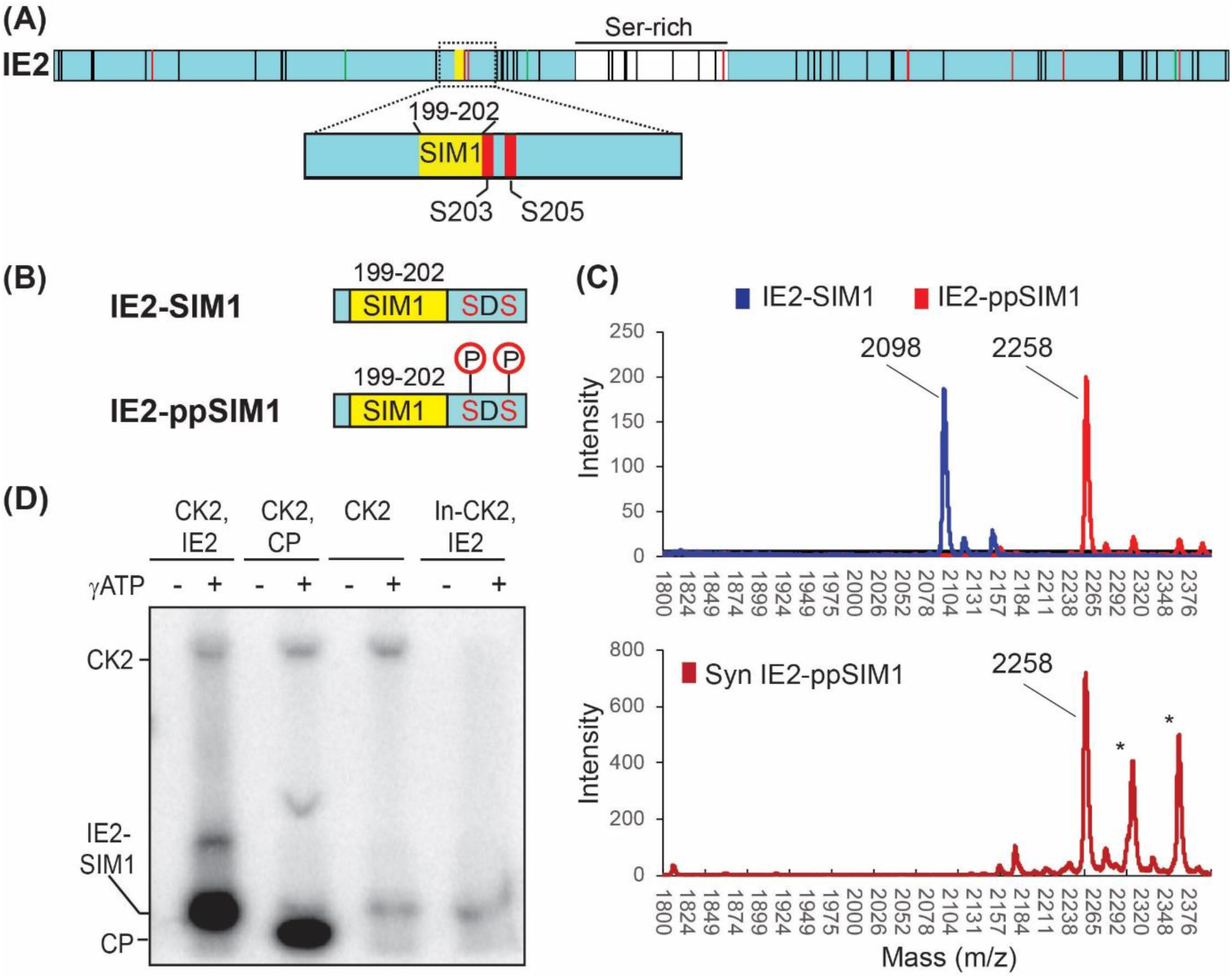
CK2 phosphorylates IE2. (A) Phosphorylation of IE2 detected in cells by Mass-spectrometry based proteomics. All vertical black lines denote detected phosphorylation sites in IE2. Green and red vertical lines denote detected phosphorylation sites in IE2, that are predicted MAPK and CK2 sites, respectively. The region around SIM1 is expanded to show that only two serines immediately adjacent to SIM1 are phosphorylated. (B) Schematic of IE2-SIM1 and IE2-ppSIM1. (C) IE2-SIM1 was incubated with CK2 and γ-ATP, run on SDS page gel and analyzed using autoradiography. CP is the control peptide that is a known substrate of CK2. In-CK2 in the last lane is inactivated CK2. The higher molecular weight band corresponds to CK2, which auto-phosphorylates itself. This band is not observed in the heat inactivated lane. (D) Mass spectra of IE2-SIM1, and IE2-SIM1 incubated with CK2 (IE2- ppSIM1). The spectra of synthesized IE2-ppSIM1 is given below as a reference. The lines with an asterisk are coming from impurities.

### Phosphorylation increases the affinity between IE2-SIM1 and SUMO1/2

Phosphorylation of SIMs may regulate the SUMO/SIM interaction (Stehmeier and Muller, 2009). NMR titrations were repeated using a synthetic peptide of IE2-SIM1 where the two serines Ser203 and Ser205 are phosphorylated (IE2-ppSIM1, Table 1), to examine the effect of Ser203,205 phosphorylation on its interaction with SUMO1/2. The pattern of CSPs observed in SUMO1 upon titration with IE2-ppSIM1 are similar to that observed during interaction with unphosphorylated IE2-SIM1 (Figure S5A), indicating that the interface of binding is identical. However, several peaks went into an intermediate exchange during the titration, suggesting that the affinity between SUMO1 and the IE2-SIM1 has increased upon phosphorylation. Indeed, the fitting of NMR chemical shifts against ligand: protein concentration yielded the dissociation constant to be 7.1 (±0.4) μM, which is 8-fold lower than unphosphorylated IE2-SIM1 (Figure S5B, Table 1). When IE2-ppSIM1 was titrated to SUMO2, the interface of interaction was similar to IE2-SIM1 (Figure S6A). The K_d_ of interaction with SUMO2 was 7.2 (±1.6) μM, confirming a significantly tighter binding upon phosphorylation (Figure S6B). In summary, phosphorylation increased the interaction between IE2-SIM1 and SUMO1/2.

It was important to determine the structure of the complex between SUMO and IE2- ppSIM1 to understand the molecular mechanism of phosphorylation induced tighter SUMO/IE2-SIM1 interaction. A ^13^C, ^15^N half-filtered NOESY-HSQC was acquired on a sample of ^13^C, ^15^N-SUMO1/IE2-ppSIM1 at the stoichiometric ratio of 1:1.5 (SUMO1:IE2- ppSIM1) (Figure 3A). ^1^H-^1^H TOCSY and ^1^H-^1^H NOESY experiments on free IE2-ppSIM1 provided the proton chemical shifts of IE2-ppSIM1. The chemical shifts of SUMO1 in the complex were assigned by comparing the ^15^N-^1^H edited HSQC and ^13^C-^1^H edited HSQC of ^13^C, ^15^N-SUMO1/IE2-ppSIM1 complex with the assignments of free SUMO1. Using the intermolecular NOEs as distance restraints; the structure of the SUMO1/IE2-ppSIM1 complex was solved by HADDOCK (Dominguez, Boelens and Bonvin, 2003) (Table 4 provides the structural statistics). Figure S7A shows the twenty lowest energy structures, which superposed well with an rmsd of 0.5Å. In the structure, the hydrophobic residues I200 and I202 packed into the hydrophobic patch between the β2 strand and α1 helix (Figure 3B). Two hydrogen bonds between D204 in IE2 and K46 in SUMO1 also stabilized the interface. The two salt-bridges between the sidechain phosphate oxygen atoms of the phosphorylated serines pSer203/pSer205 in IE2-ppSIM1 and K39 of the β2-strand in SUMO1 is responsible for the higher affinity between IE2-ppSIM1 and SUMO1 (Figure 3C).

**Figure 3.**
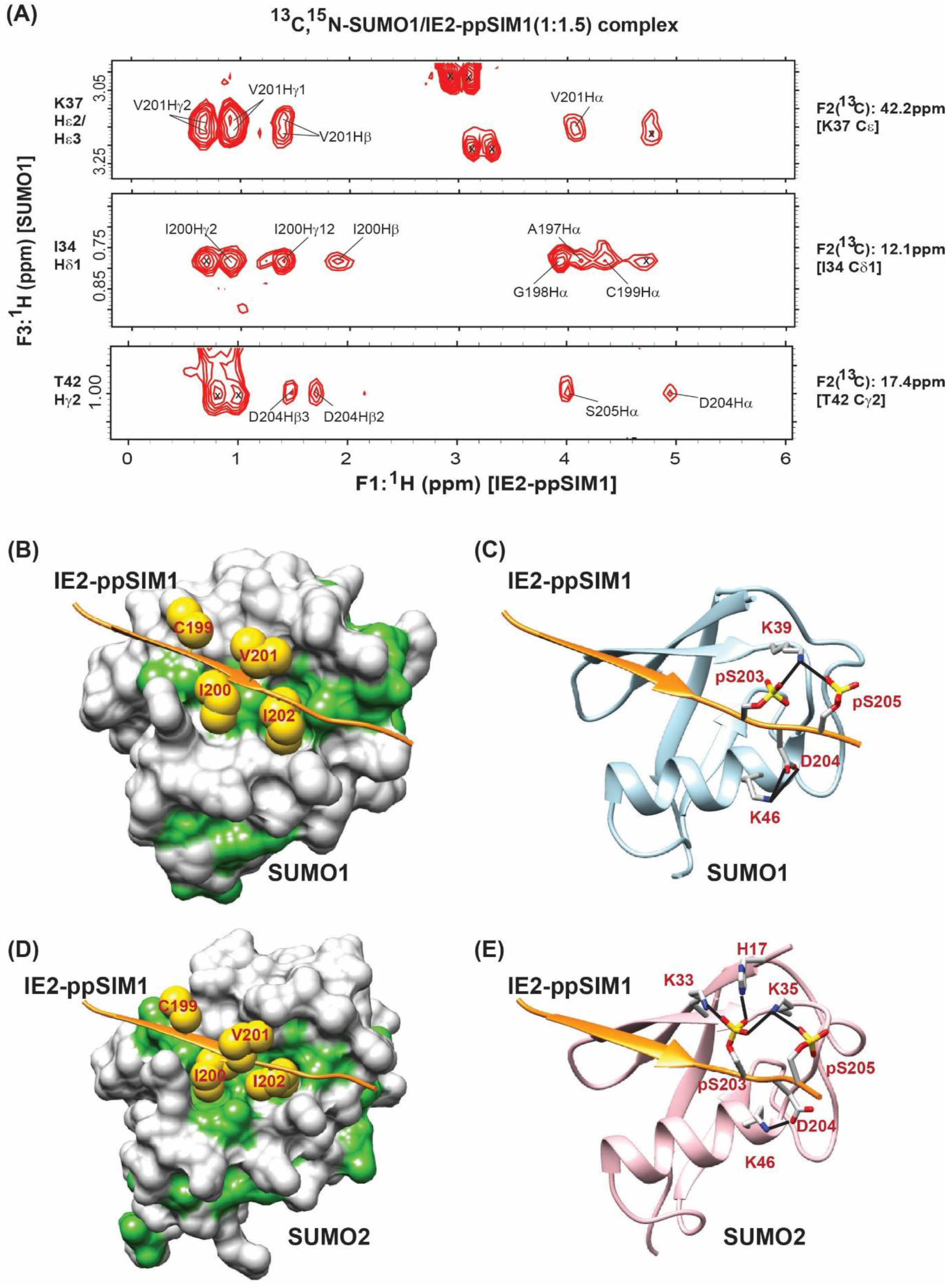
Molecular basis of the enhanced interaction between SUMO and IE2- ppSIM1. (A) Selected strips from the ^13^C, ^15^N half-filtered NOESY spectra depicting intermolecular NOEs between ^13^C-bonded protons of ^13^C, ^15^N-labeled SUMO1, and unlabeled IE2-ppSIM1. ^13^C and ^1^H assignment of SUMO1 atoms are given on the right and left of the strips, respectively. The protons of IE2-ppSIM1 that show NOEs to SUMO1 are assigned. (B) and (D) highlights the hydrophobic interactions in the SUMO1/IE2-ppSIM1 and SUMO2/IE2-ppSIM1 complexes, respectively. The SUMO1/2 surface is colored white, except the hydrophobic patches are colored green. The IE2-ppSIM1 backbone is shown as an orange ribbon. The sidechains of central hydrophobic residues ‘CIVI’ are shown as yellow spheres. (C) and (E) shows the hydrogen bonds between phosphorylated side-chains of IE2-SIM1 with SUMO1 and SUMO2, respectively. The hydrogen bonds are shown as black lines. The two phosphoserines and the residues in SUMO1/2 that form hydrogen bonds are shown. Nitrogen atoms are colored in blue, oxygen atoms are colored in red, and phosphorous atoms are colored in yellow.

**Table 4.**
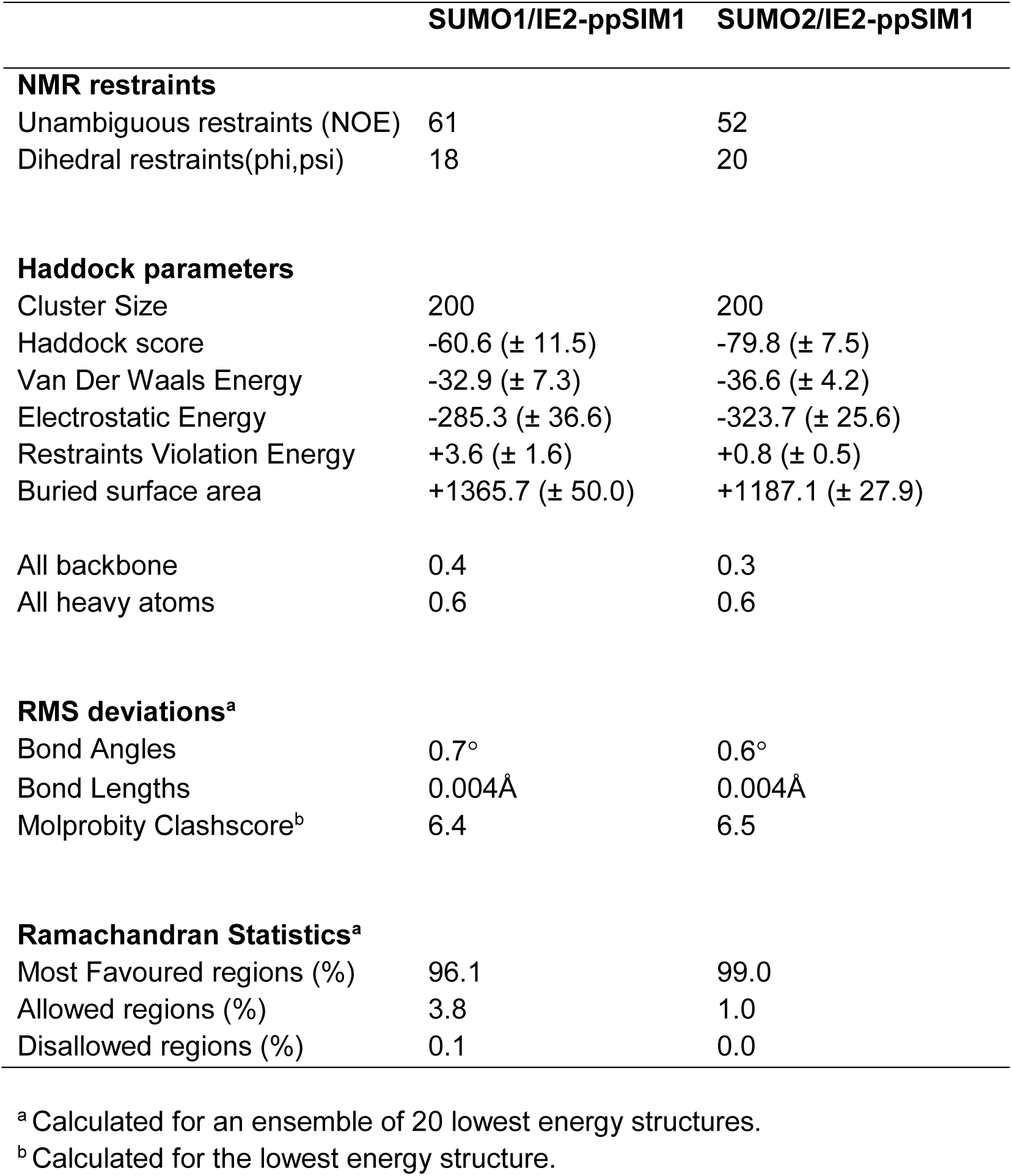
NMR and refinement statistics of the SUMO/IE2-ppSIM1 complexes.

|  | **SUMO1/IE2-ppSIM1** | **SUMO2/IE2-ppSIM1** |
| --- | --- | --- |
| **NMR restraints** |  |  |
| Unambiguous restraints (NOE) | 61 | 52 |
| Dihedral restraints(phi,psi) | 18 | 20 |
| **Haddock parameters** |  |  |
| Cluster Size | 200 | 200 |
| Haddock score | -60.6 (± 11.5) | -79.8 (± 7.5) |
| Van Der Waals Energy | -32.9 (± 7.3) | -36.6 (± 4.2) |
| Electrostatic Energy | -285.3 (± 36.6) | -323.7 (± 25.6) |
| Restraints Violation Energy | +3.6 (± 1.6) | +0.8 (± 0.5) |
| Buried surface area | +1365.7 (± 50.0) | +1187.1 (± 27.9) |
| All backbone | 0.4 | 0.3 |
| All heavy atoms | 0.6 | 0.6 |
| **RMS deviations<sup>a</sup>** |  |  |
| Bond Angles | 0.7° | 0.6° |
| Bond Lengths | 0.004Å | 0.004Å |
| Molprobity Clashscore<sup>b</sup> | 6.4 | 6.5 |
| **Ramachandran Statistics<sup>a</sup>** |  |  |
| Most Favoured regions (%) | 96.1 | 99.0 |
| Allowed regions (%) | 3.8 | 1.0 |
| Disallowed regions (%) | 0.1 | 0.0 |

<sup>a</sup> Calculated for an ensemble of 20 lowest energy structures.
<sup>b</sup> Calculated for the lowest energy structure.

The structure of the SUMO2/IE2-ppSIM1 complex was studied to determine if a similar mechanism operates between phosphorylated IE2-SIM1 and SUMO2. We prepared a sample of ^13^C, ^15^N-labeled SUMO2/IE2-ppSIM1 at the stoichiometric ratio of 1:1.5 (SUMO2:IE2-ppSIM1). Half-filtered ^13^C, ^15^N NOESY-HSQC of this sample detected several intermolecular NOES between SUMO2 and IE2-ppSIM1 (Figure S7B), which were used to determine the SUMO2/IE2-ppSIM1 structure. The twenty lowest energy structures superimposed with a low rmsd of 0.4Å (Figure S7C). Similar to SUMO1, the IE1-I200 and I202 sidechain are buried in the hydrophobic patch between β2 and α1 of SUMO2 (Figure 3D). The oxygen atom of phosphate groups of pSer203 and pSer205 form salt-bridges with K33, K35 in β2 and H17 in β1 (Figure 3E). The additional salt-bridges in the between the phosphoserines in IE2-ppSIM1 and SUMO, explain the higher affinity of the IE2/SUMO non-covalent interaction upon phosphorylation of IE2.

### IE2-SIM1 enhanced SUMOylation of IE2

SIMs often regulate the SUMOylation of a protein in-cis (Kolesar *et al*., 2012). The identified SUMOylation sites in IE2 are K175 and K180, which are adjacent to IE2-SIM1. *In-vitro* SUMOylation assays were carried out to assess the effect of IE2-SIM1 on the SUMOylation of IE2. A FITC was attached to the N-terminal region of IE2 (aa: 172-210), which we termed as IE2 N-Terminal Domain (IE2-NTD, Figure 4A). IE2-NTD includes both the SUMOylation sites K175 and K180 as well as the IE2-SIM1. Sumoylation reactions were carried out using IE2-NTD as the substrate. Robust and rapid poly-SUMOylation of IE2-NTD with SUMO1 could be observed in the reaction (Figure 4B). The reaction was repeated with a SIM mutated IE2-NTD (IE2-NTDm), where the central hydrophobic residues CIVI were mutated to AAAA. IE2-NTDm showed a significantly reduced rate of SUMOylation (Figure 4C). The efficiency of SUMOylation was estimated by the decay rate of unmodified or apo IE2-NTD signal against time. Faster decay reflected a higher rate of SUMOylation. Apo IE2-NTD signal decayed rapidly compared to IE2-NTDm, which indicated efficient SUMOylation of IE2-NTD due to the presence of IE2-SIM1 (Table 2, Figure 4D). When SUMOylation of IE2- NTD was repeated with SUMO2, the rate of SUMOylation was again higher in IE2-NTD than IE2-NTDm (Figure 4E, Figure S8A, S8B and Table 2), indicating that the IE2-SIM1 promotes IE2 SUMOylation.

**Figure 4.**
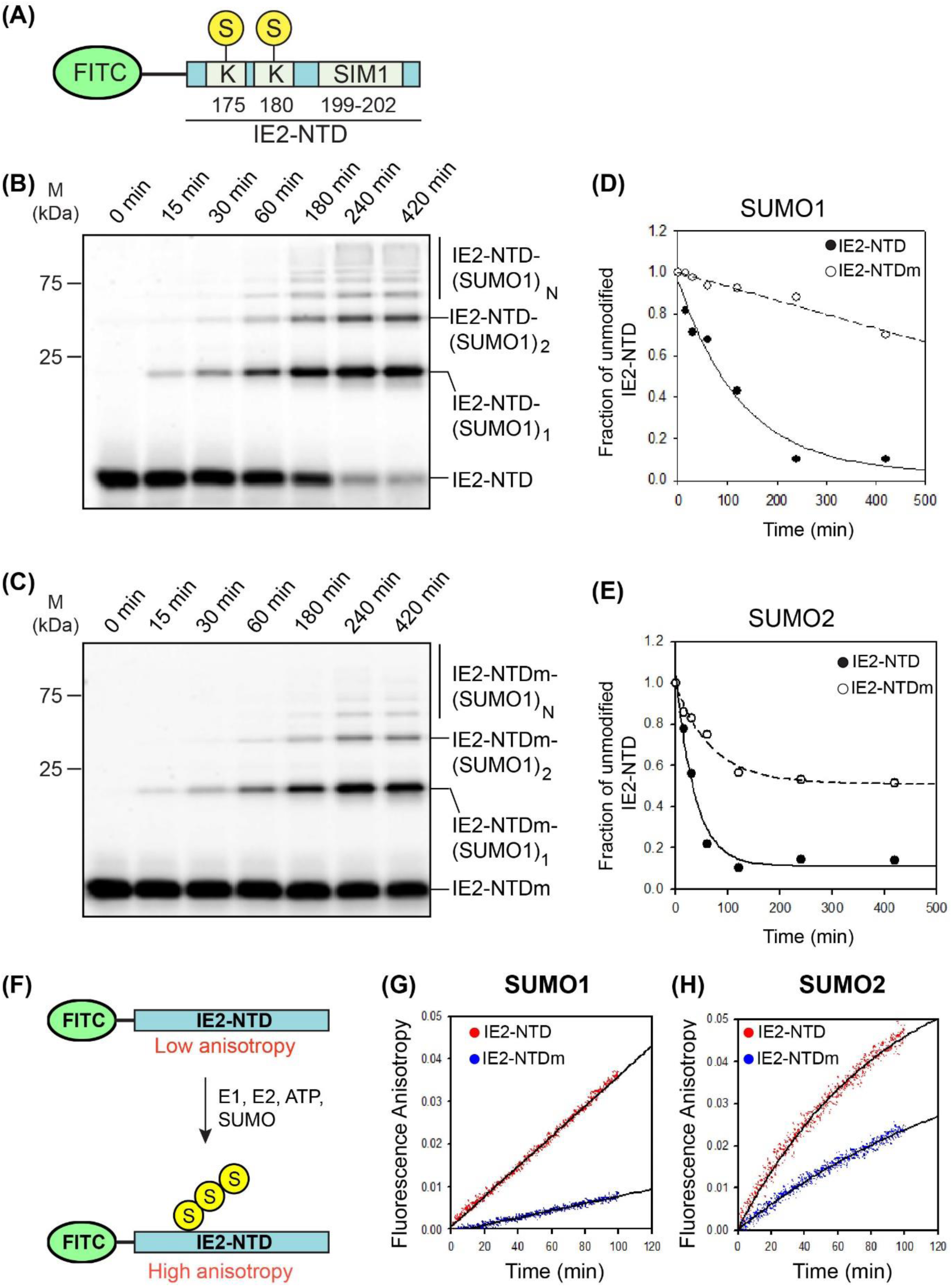
*In-vitro* SUMOylation of IE2-NTD. (A) FITC fluorophore-labeled IE2-NTD used as a substrate in SUMOylation assays. The substrate lysines and SIM1 are shown. (B) The products of the SUMOylation reaction with SUMO1 and IE2-NTD as the substrate is resolved on the SDS-PAGE gel and imaged with a filter at 519 nm corresponding to FITC fluorescence. Bands of free IE2-NTD or conjugated with one, two or multiple (n) SUMO1 are marked. The time-points are given on the top of the gel. (C) Same as (B) except that IE2- NTDm (CIVI to AAAA) was used as the substrate. (D) and (E) Fraction of free IE2-NTD is plotted against time for SUMOylation reactions using SUMO1 and SUMO2, respectively. (F) Experimental design to monitor SUMOylation of IE2-NTD in real-time. (G) Change of fluorescence anisotropy with time for IE2-NTD and IE2-NTDm in a SUMOylation reaction using SUMO1. (H) The same as in (G) using SUMO2 instead of SUMO1.

**Table 2.**
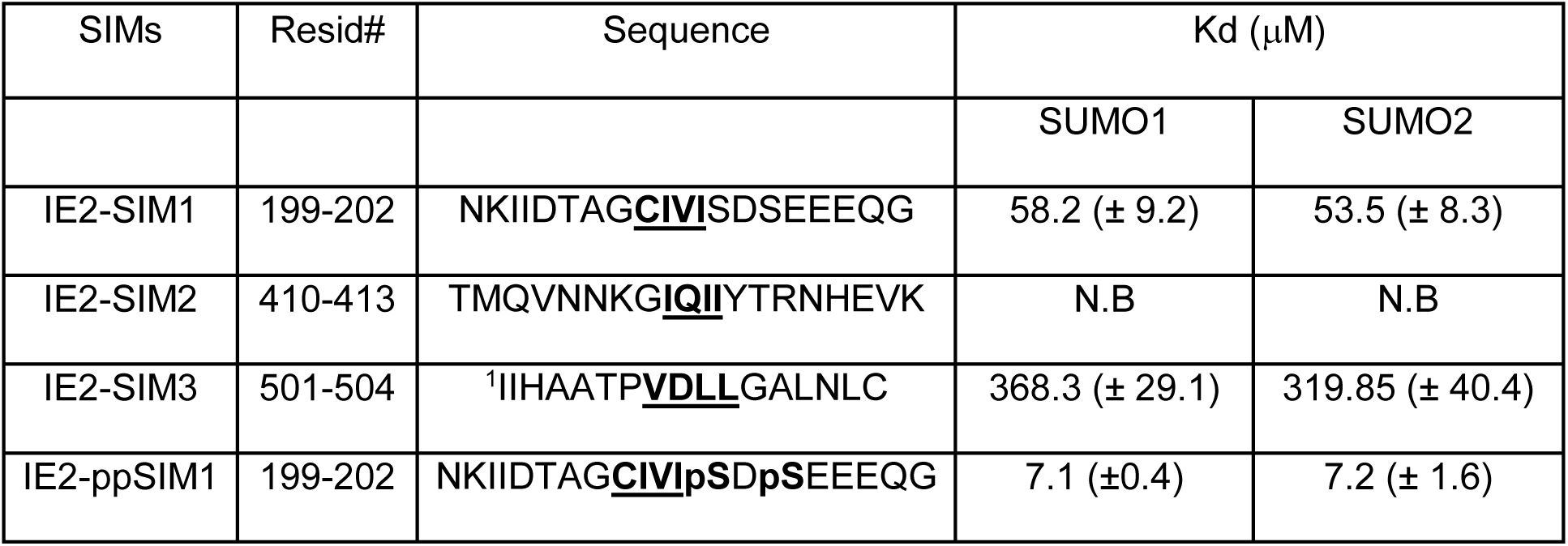
The rate of SUMOylation in IE2-NTD

The poly-SUMOylation of IE2-NTD would increase its size and anisotropy. Hence, the SUMOylation of the substrate IE2-NTD can be monitored in real time by measuring the change in fluorescence anisotropy of IE2-NTD (Figure 4F). As a control, a reaction without ATP was carried out, where the absence of SUMOylation did not change the anisotropy (Figure S8C). In the SUMOylation reaction using SUMO1 and IE2-NTD/IE2-NTDm as the substrate, the rate of increase in fluorescence anisotropy of was higher for IE2-NTD than IE2-NTDm, confirming that the IE2-SIM1 increased SUMOylation of IE2-NTD. When the reaction was repeated with SUMO2, the rate of SUMOylation was higher than SUMO1 (Table 2). This could be because SUMO2 has a consensus SUMOylation motif at K11, which enhances the rate of poly-SUMOylation. Nevertheless, IE2-SIM1 increased the rate of SUMOylation (Figure 4G and Table 2). Together, the IE2-SIM1 enhanced the SUMOylation of IE2 significantly.

### Mechanism of SIM enhanced IE2 SUMOylation

In principle, IE2-SIM1 could increase the SUMOylation of IE2 either by reducing K_m_ (increasing affinity) between the substrate IE2 and the enzyme UBC9∼SUMO conjugate or by increasing the activity (K_cat_) of the enzyme UBC9∼SUMO. To determine the exact effect of IE2-SIM1, we carried out a kinetic study of the IE2 SUMOylation. IE2-NTD is poly-SUMOylated rapidly, which made quantification of the SUMOylated species difficult. We used the mutant of SUMO2 (K11R-SUMO2) that is deficient in poly-SUMOylation, which allowed us to effectively monitor the mono-SUMOylated IE2 (Figure 5A, Figure S9). Kinetic analysis of IE2 SUMOylation revealed that IE2-SIM1 reduced the K_m_ of IE2 by 2.5 fold (Figure 5B), but does not alter the V_max_ (K_cat_) significantly (Figure 5C). Thereby, IE2 SIM1 increases specificity constant (K_cat_/K_m_) by decreasing K_m_ of the SUMOylation reaction. (Table 3). The presence of IE2-SIM1 significantly increased the SUMOylation of IE2 in cellular conditions (Figure 5D, 5E).

**Figure 5.**
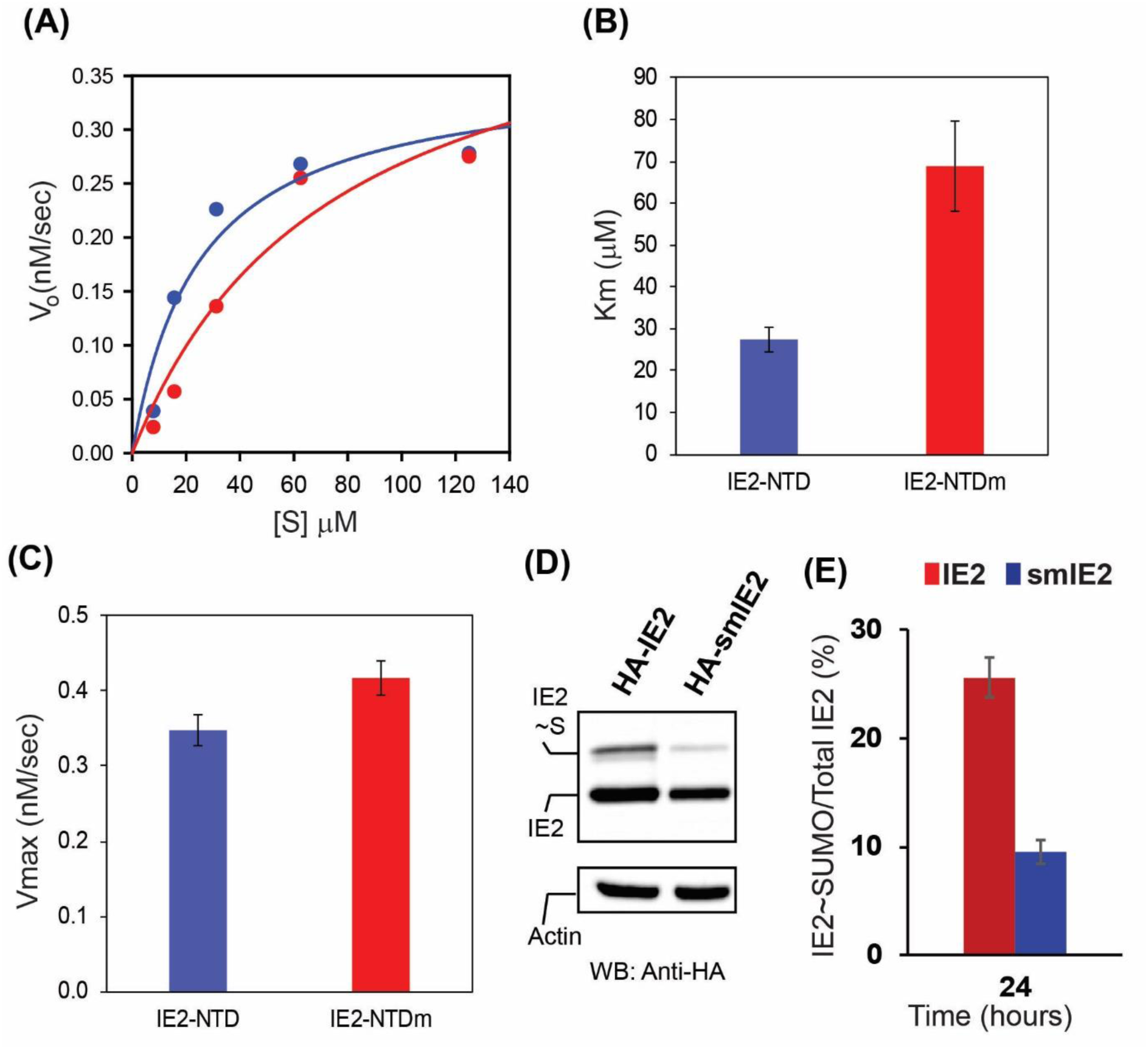
The IE2-SIM1 enhances SUMOylation of IE2. (A) Kinetic data for SUMOylation of IE2-NTD and IE2-NTDm. (B) and (C) are the calculated K_m_ and V_max_, respectively. (D) IE2 and SIM mutant IE2 (smIE2) SUMOylation detected in HEK293T cells. (E) The fraction of IE2∼SUMO over total IE2 is quantified from (D) and plotted.

**Table 3.** Kinetic Analysis of IE2 SUMOylation

| Substrate | $V_{\max}$ (nM/s) | $K_{\text{cat}}$ (1/s) | $K_m$ ($\mu\text{M}$) | Specificity,<br>$K_{\text{cat}}/K_m$<br>($\mu\text{M}^{-1} \text{s}^{-1}$) | Temp<br>($^{\circ}\text{C}$) |
| --- | --- | --- | --- | --- | --- |
| IE2-NTD | 0.35 ($\pm 0.04$) | $8.7 \times 10^{-3}$ ($\pm 5.2 \times 10^{-4}$) | 27.5 ($\pm 3.0$) | $3.2 \times 10^{-4}$ | 25 |
| IE2-NTDm | 0.42 ($\pm 0.04$) | $10.4 \times 10^{-3}$ ($\pm 5.6 \times 10^{-4}$) | 68.9 ($\pm 10.7$) | $1.5 \times 10^{-4}$ | 25 |
| IE2-NTD | 0.12 ($\pm 0.01$) | $3.3 \times 10^{-3}$ ($\pm 1.2 \times 10^{-4}$) | 22.1 ($\pm 2.9$) | $1.4 \times 10^{-4}$ | 16 |
| IE2-ppNTD | 0.16 ($\pm 0.01$) | $3.9 \times 10^{-3}$ ($\pm 0.5 \times 10^{-4}$) | 6.7 ($\pm 0.4$) | $6.8 \times 10^{-4}$ | 16 |

Three mechanisms of IE2-SIM1 enhanced SUMOylation are possible (Figure 6A-C). The binding of IE2-SIM1 to the SUMO conjugated at the active site of Ubc9 can increase the rate of SUMOylation (Figure 6A). Alternately, SUMO has a non-covalent interaction with UBC9 at its “backside binding” area, and this interaction could also enhance IE2 SUMOylation (Figure 6B). Moreover, UBC9 is SUMOylated covalently at K14, which could also impact IE2-SUMOylation (Figure 6C). NMR titration experiments studied the hypothesis in Figure 6A. ^15^N-labeled wt-UBC9 was titrated with IE2-NTD, and the observed CSPs in UBC9 are plotted in Figure 6D. The IE2-NTD was truncated from the N-terminal end such that K180 was the sole acceptor lysine present in IE2-NTD. The high CSPs at the α’-α2 and α2-α3 loops indicated the binding site of acceptor K180 and UBC9. The other significant CSPs were observed in the α1-β1 region. The pattern of CSPs was consistent with NMR titrations of UBC9 with SUMO-acceptor sites of p53 and c-Jun (Lin *et al*., 2014). Then, C93K-UBC9∼^15^N-SUMO1 conjugates were purified and titrated with IE2-NTD. The pattern of CSPs on SUMO1 matched with the CSPs observed in isolated SUMO1/IE2-SIM1 complex (Figure 6E), indicating that the IE2-SIM1 within the IE2-NTD binds to the same interface on the SUMO1 conjugated to UBC9. IE2-NTD does not bind to the UBC9 active site when the active site is mutated to lysine (C93K-UBC9), and hence the binding to the C93K-UBC9∼^15^N-SUMO1 conjugate could not be studied (Figure S10A). Nevertheless, combining the NMR data of IE2-SIM1/SUMO interaction, IE2-NTD/UBC9∼SUMO1 interaction and the UBC9∼SUMO1 structure (PDB: 1Z5S), a model of UBC9∼SUMO1/IE2-NTD complex was determined by Xplor-NIH (Figure 6F and Figure S11A). The structure shows how the IE2- SIM1 binds to SUMO1, while the acceptor lysine K180 attacks the active site (Figure 6I). Interestingly, the essential negatively charged residue of the ψKx<u>D/E</u> motif, which is E182 in this context, forms a salt-bridge with the positively charged K101 in the β4α2 loop of UBC9. K101 is essential for the recognition of substrates like p53, PML and IκB (Tatham, Chen and Hay, 2003). The molecular details provided by the structural model explain the underlying mechanism of substrate recognition by K101 in UBC9.

**Figure 6.**
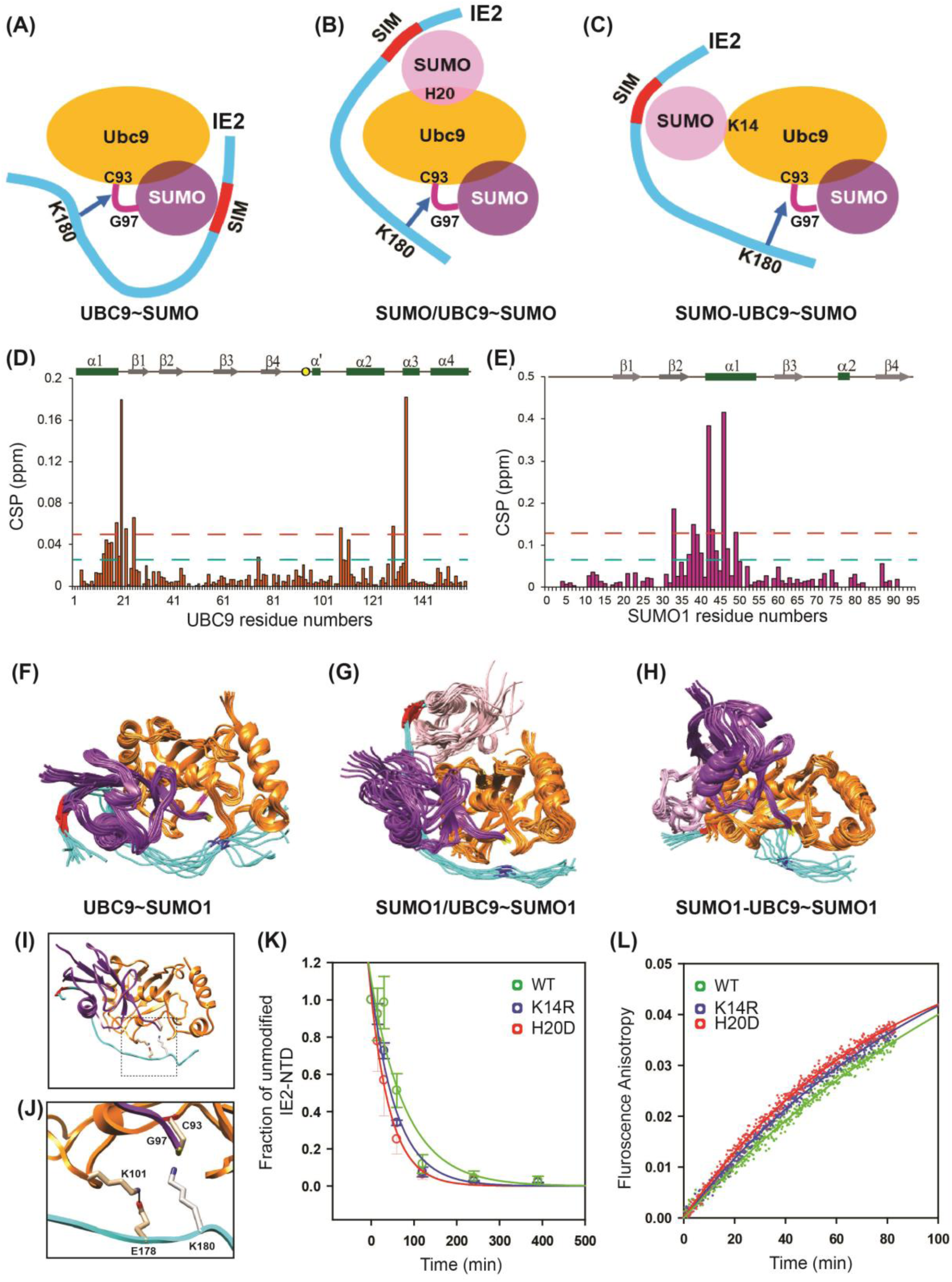
Mechanism of cis SUMO-E3 ligase activity of IE2. The three possible mechanisms are shown in (A), (B) and (C). C93 is the catalytic cysteine in UBC9, which is conjugated to G97 in SUMO1. The critical residue H20 for non-covalent UBC9/SUMO interaction is shown. The critical residue K14 for covalent UBC9/SUMO interaction is also shown. (D) The CSPs observed in UBC9 upon titration with IE2-NTD. (E) The CSPs in SUMO1 within the UBC9∼SUMO1 conjugate, upon titration with IE2-NTD. (F) The ten lowest energy model structures of the UBC9∼SUMO1/IE2-NTD complex. UBC9, SUMO1, and IE2-NTD are color-coded as in (A). The active site C93 is colored purple, G97 of SUMO1 is colored yellow and K180 is colored blue. (G) Same as in (F) for the SUMO1/UBC9∼SUMO1/IE2-NTD, where SUMO1 is non-covalently bound to UBC9. (H) Same as in (F) for the SUMO1- UBC9∼SUMO1/IE2-NTD complex, where SUMO1-UBC9 denotes the SUMO1 covalently linked to K14 of UBC9. (I) Lowest energy structure of the UBC9∼SUMO1/IE2-NTD complex. (J) A close-up of the active site shows that E178 forms a salt-bridge with K101, while K180 attacks the active site. The inset is shown the complete structure, where the zoomed region is marked with a box. (K) Gel shift assay and (L) fluorescent anisotropy assay monitored the rate of SUMOylation using either wt, K14R or H20D UBC9.

A structural model corresponding to Figure 6B was build using the NMR CSP data, the structure of UBC9∼SUMO1 and the structure of UBC9/SUMO (PDB), where SUMO binds to the ‘backside’ beta-sheet of UBC9 (Figure 6G, S11B and S12). Again, the E178 forms a salt-bridge with K101. The structural model of Figure 6C was determined using covalently linked SUMO1-UBC9 structure (PDB ID: 2VRR, Figure 6H, S11C, and S12). In this case, the E178 prefers to form a salt-bridge with K74, which is present on the beta-strand β4. K74 is vital to identify substrates like RANGAP1 (Bernier-Villamor *et al*., 2002). K175 is further away from the IE2-SIM1 than K180 and had more flexibility to access the active site of UBC9 when the IE2-SIM1 binds to either conjugated, covalently linked or non-covalently linked SUMO. Overall, structural modeling suggested that all the three possibilities of SIM enhanced SUMOylation are possible in IE2-NTD.

SUMOylation assays were performed with appropriate substitutions of UBC9 to delineate the effects of each mechanism. The non-covalent interaction between UBC9 and SUMO involves Histidine 20 in UBC9, and the H20D mutation abolishes the interaction (Knipscheer *et al*., 2007). Alternately, The K14R UBC9 abolishes the covalent SUMO conjugation. The IE2-NTD SUMOylation was studied by initiating a SUMOylation reaction using E1, SUMO1, and wt-UBC9 or H20D-UBC9 or K14R-UBC9. Gel-shift assays or real-time fluorescence anisotropy assay monitored the SUMOylation of IE2-NTD. When the covalent or non-covalent mediated interactions between UBC9 and SUMO were abolished individually, the rate of IE2-NTD SUMOylation did not change significantly (Figure 6H, 6I). A possible cause of the result is that the mechanisms involving the covalent or non-covalent UBC9/SUMO interaction do not contribute much to IE2 SUMOylation. However, the three mechanisms could be redundant, such that when one is interrupted, the others compensate.

### Phosphorylation of IE2-SIM1 further enhances IE2 SUMOylation

The SUMOylation assays were repeated with IE2-NTD and phosphorylated IE2-NTD (IE2- ppNTD) to find out whether the phosphorylation of IE2-SIM1 enhances the SUMOylation IE2. FITC-tagged IE2-NTD was incubated with CK2 for phosphorylation. The CK2 was subsequently inactivated, and the IE2-NTD was checked by Mass-spec to ensure that the serines Ser203 and Ser205 were phosphorylated. The phosphorylated IE2-NTD (IE2- ppNTD) was later used as a substrate in the SUMOylation assay (Figure 7A). The rate of SUMOylation was higher in IE2-ppNTD than IE2-NTD (Figure 7B). At 60min, 50% of total IE2-ppNTD was SUMOylated, while IE2-NTD was only SUMOylated up to 30% (Figure 7C).

**Figure 7.**
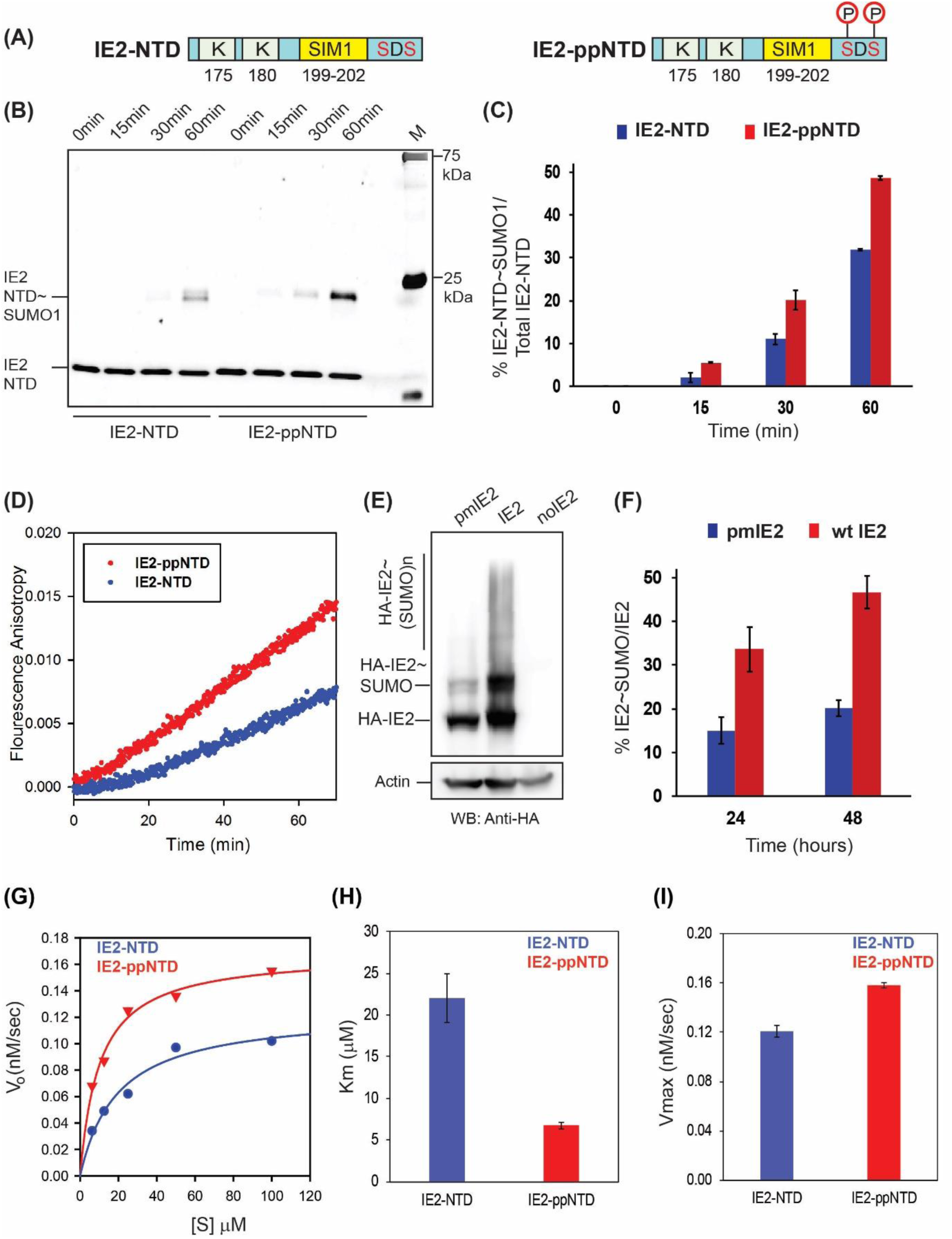
Phosphorylation enhanced SUMOylation of IE2-NTD. (A) Schematic of IE2-NTD and the phosphorylated IE2-NTD. (B) SUMOylation of IE2-NTD and IE2-ppNTD against time. The SUMOylation reaction using either IE2-NTD or IE2-ppNTD was run for different times, resolved on the SDS-PAGE gel and imaged with a filter at 519 nm corresponding to FITC fluorescence. The time of reaction is given on the top. M stands for the marker. (C) The IE2-NTD∼SUMO conjugate was quantified and plotted against time for IE2-NTD and IE2-ppNTD. (D) Real-time fluorescence anisotropy measurement of IE2-NTD and IE2- ppNTD SUMOylation. (E) The SUMOylation of IE2 and phosphorylation mutant pmIE2 observed in HEK293T cells. Cells lysates 48 hours post-transfection were separated on SDS page and blotted with anti-HA. (F) The ratio of conjugated and total IE2 is quantified from (E) and plotted against the time of transfection. (G) Michelis Menten curves for IE2-NTD and IE2-ppNTD. (H) and (I) are the calculated K_m_ and V_max_, respectively.

The altered rate of SUMOylation upon SIM phosphorylation was also monitored in real-time by the fluorescence anisotropy assay using SUMO1 and IE2-NTD/IE2-ppNTD as the substrate (Figure 7D). The rate of increase in anisotropy was measured to be 1.2 x 10^-4^ /min for wt IE2-NTD, which almost doubled to 2.1 x 10^-4^ /min for IE2-ppNTD, indicating that phosphorylation of IE2-SIM1 enhances the SUMOylation of IE2-NTD.

SUMOylation of IE2 was measured in HEK293T cells to assess if the effect of phosphorylation on SUMOylation of IE2 is persistent in full-length IE2 in cellular conditions. HA-tagged IE2 or a double phospho-inactive mutant IE2pm: Ser203A/Ser205A was co-transfected with SUMO1, lysed 24/48 hours post-transfection and probed with HA antibody (Figure 7E, F). The amount of IE2pm∼SUMO1 was significantly lower than IE2∼SUMO1, indicating that the phosphorylation of serines near SIM1 enhances SUMOylation of IE2. The enzyme kinetics experiments were repeated with IE2-NTD and IE2-ppNTD. The experiments were carried out at 16°C to slow down the reaction and effectively measure the kinetics (Figure 7G, Figure S13). The kinetic analysis given in Table 3, indicates that the phosphorylation reduces the Km of the reaction by 3.5-fold but does not change the V_max_ significantly (Figure 7H, 7I). Together, phosphorylation of two serines near IE2-SIM1 by CK2 increases the SUMOylation of IE2.

### Phosphorylation of IE2-SIM1 increases transactivation by IE2

IE2 functions as a transactivator for various viral promotors and auto-repressor for its promoter. IE2 SUMOylation and IE2 SIM1 are essential for its function. Ser203, Ser205 phosphorylation enhanced both the IE2-SIM1/SUMO interaction and IE2-SUMOylation. The role of phosphorylation on the IE2 mediated transactivation was examined by a luciferase assay, where the luciferase gene is expressed under IE2 responsive promoter (pUL54-Luc). Consequently, the level of luciferase expression was proportional to the IE2 transactivation activity. A phosphorylation inactive S203A, S205A mutant (pmIE2) and a SIM mutant (CIVI to AAAA, smIE2) of IE2 were designed. PUL54-Luc was transfected into HEK293T along with HA-IE2 (wt or mutants) and SUMO1 (Figure S14). Transactivation activity of pmIE2 is reduced by ∼25% in comparison to wt-IE2, indicating that phosphorylation of Ser203 and Ser205 is vital for transactivation (Figure 8A). Interestingly, the activity of pmIE2 is comparable to smIE2, indicating that SIM phosphorylation is necessary for the SUMO-dependent transactivation activity of IE2.

**Figure 8.**
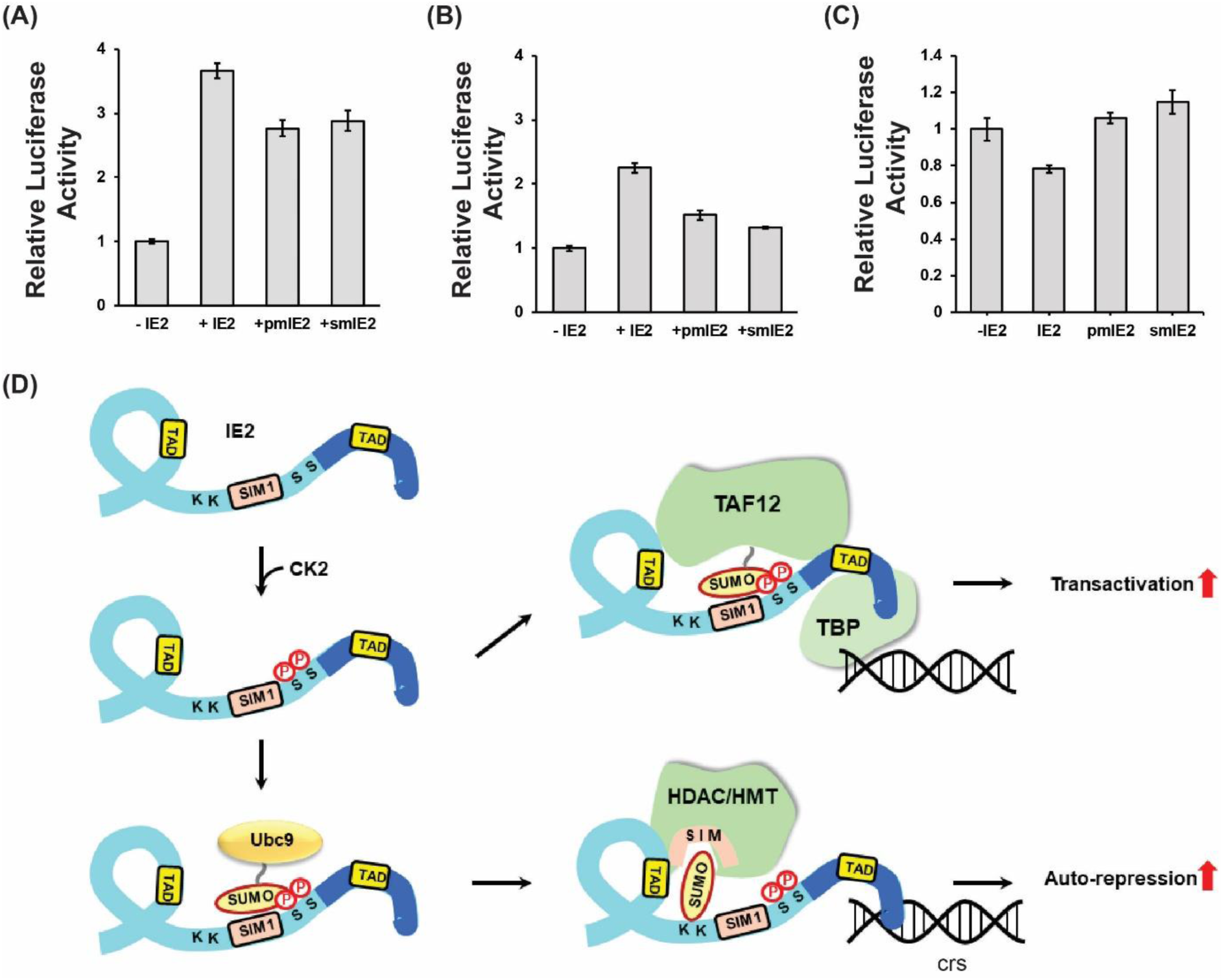
Functional implications of phosphorylation induced enhanced SIM/SUMO interaction and SUMOylation. (A) Luciferase transactivation assays were performed in HEK293T cells 36 hours post-transfection with or without IE2 or mutants of IE2, and SUMO1. The relative Luciferase activity is plotted against the IE2 or its mutants. (B) The same is repeated without transfection of SUMO1. (C) Luciferase auto-repression activity was monitored using IE2 and its mutants. C denotes the control where IE2 was not transfected. (D) Model of phosphorylation induced enhanced IE2 transactivation/auto-repression activity. IE2 is colored in light-blue, and the DNA binding domain (DBD) is colored in dark blue. IE2 binds to the TATA-box binding protein (TBP), which binds to the promoter. Phosphorylation increases the interaction between IE2-SIM1 and SUMOylated transcription factors (e.g., TAF12) to enhance the transactivation activity. IE2 can directly bind to the cis-regulatory sequences (crs) for auto-repression via the DBD. Phosphorylation can enhance SUMOylation of IE2 to increase its association with the chromatin modifiers like HDAC/HMT/CoREST complex and increase auto-repression. HDAC: Histone Deacetylase and HMT: Histone Methyltransferase.

The effect of phosphorylation was also checked with co-transfection of only IE2 and pUL54-Luc, but not SUMO. Interestingly, the activity of wt-IE2 with endogenous SUMO reduced by 50% in comparison to overexpressed SUMO (Figure 8B). IE2 SUMOylation increases upon overexpression of SUMO (Figure S15). The increase of both SUMOylation and transactivation activity of IE2 upon overexpression of SUMO underscores the importance of IE2 SUMOylation for its transactivation activity. Alternately, increased SUMOylation of IE2 associated transcription factors may also increase the IE2 mediated transactivation (Figure 8D). Nevertheless, the activity of pmIE2 dropped by 40% in comparison to wt-IE2, emphasizing the significance of SIM phosphorylation when the amount of SUMOylated transcription factors or SUMO is not abundant (Figure 8B). Additionally, the importance of Ser203, 205 phosphorylation on the auto-repression activity of IE2 was studied using similar luciferase assays where Luc was expressed under MIEP. Expression of luciferase decreases on over-expression of IE2 indicating MIEP repression by IE2. Interestingly, MIEP repression is relieved in pmIE2, indicating the importance of SIM phosphorylation for the auto-repression activity (Figure 8C). IE2 SUMOylation is important for MIEP repression and SIM phosphorylation enhances IE2 SUMOylation. Our data suggest that SIM phosphorylation augments SUMOylation dependent activity of IE2 as an auto-repressor.

## Discussion

SUMOylation is an essential component of cell signaling pathways, and it is unsurprising that the intracellular viruses have co-evolved to exploit the host cell SUMOylation system (Wimmer, Schreiner and Dobner, 2011; Everett, Boutell and Hale, 2013). However, very little is known about the molecular mechanism of how viruses co-opt the machinery. For example, it is known that IE2 binds SUMO non-covalently and this interaction is indispensable for SUMOylation of IE2 (Berndt *et al*., 2009). However, the molecular details of IE2/SUMO non-covalent interaction and its role in the SUMOylation of IE2 are unknown. In this work, NMR titrations studies determined that apart from the known N-terminal SIM, IE2 includes another C-terminal SIM. Unlike some other SIMs that have paralog specificity, IE2-SIMs binds equally well to both SUMO1 and SUMO2. The C-terminal SIM is located in the DNA-binding domain of IE2, and the non-covalent interaction with SUMO may affect its binding to the cis-regulatory elements in MIEP. The N-terminal SIM is located near the SUMOylation sites and is indispensable for the SUMOylation of IE2. The binding of N-terminal SIM to SUMO is 8- fold tighter than the C-terminal SIM.

The structural studies showed IE2-SIM1 binds to the β2α1 groove of SUMO and forms a parallel beta-strand with β2. The four central hydrophobic residues in IE2-SIM ^199^CIVI^202^ pack against the hydrophobic interface between β2 and α1. SUMOylation monitored by fluorescence spectroscopy revealed that the IE2-SIM1/SUMO interaction enhances the rate of SUMOylation. The effect is more for SUMO1 than SUMO2, probably because the inherent SUMOylation site in SUMO2 makes the reaction more processive. Kinetic studies indicated that IE2-SIM1 increases the affinity between the substrate (IE2) and enzyme complex (UBC9∼SUMO), but does not change K_cat_ of the SUMOylation reaction. Typically, E3s decrease the binding constants of E2∼Ubl for substrate (K_m_) and increase the turnover rate of E2 (K_cat_) to increase the specificity constant (K_cat_/K_m_) of the reaction. In SUMOylation, thioester conjugation of SUMO to UBC9 reduces the flexibility of residues Cys93, Asp127, and Pro128, which help the acceptor lysine of the substrate to attack the active site (Tozluoǧlu *et al*., 2010). Consequently, UBC9∼SUMO active site is constitutively primed for SUMOylation, and reducing the K_m_ is sufficient to induce catalysis. IE2 can be considered as a SUMO cis-E3, where the IE2-SIM1 binds to SUMO in the UBC9∼SUMO enzyme complex to bring the complex in the vicinity of SUMOylation sites K175/K180 and increase the rate of SUMOylation.

The thioester conjugated SUMO, covalent bound SUMO, and non-covalent bound SUMO can potentially interact with IE2-SIM1 to enhance the rate of SUMOylation. Generally, the mechanism used by the enzyme/substrate complex depends on the substrate. For example, although the presence of SIM enhanced SUMOylation of SP100, Daxx, PML and TDG, covalent binding of SUMO to UBC9 only enhanced SUMOylation of SP100 and Daxx, but not of PML and TDG (Knipscheer *et al*., 2008). In the case of IE2, modeling studies indicated that sterically all three mechanisms are possible. However, disruption of either covalent or non-covalent SUMO interaction had little effect on the rate of SUMOylation. The thioester conjugated UBC∼SUMO/IE2-NTD complex could be energetically favorable and the only mechanism at play here. Alternately, all three mechanisms could be redundant for IE2.

*In vitro*, CK2 phosphorylates IE2 at the Serine-rich segment and the region fragment 5B (aa: 180-252) (Barrasa, Harel and Alwine, 2005). However, the exact sites of phosphorylation in fragment 5B are unknown. The putative CK2 phosphorylation sites in fragment 5B are Ser203 and Ser205. We report that in cellular conditions, Ser203 and Ser205 are indeed phosphorylated in IE2. The same residues are also phosphorylated by CK2 *in-vitro*. These serines are immediately next to IE2-SIM1 and influence both the covalent and non-covalent binding between SUMO and IE2. The affinity of IE2-SIM1/SUMO non-covalent interaction increases significantly by 8-fold upon phosphorylation of the adjacent serines. Structures of SUMO/IE2-ppSIM1 complexes show that pSer203 and pSer205 form additional salt bridges with positively charged residues in SUMO to tighten the interaction between SIM and SUMO. The stronger non-covalent interaction also impacts the covalent interaction between SUMO and IE2. *In-vitro* SUMOylation of IE2 increases significantly upon phosphorylation of Ser203 and Ser205. Kinetic analysis revealed that the phosphorylated SIM reduces the K_m_ between UBC9∼SUMO and IE2 to increase the specificity of the reaction by three-fold. Moreover, in cellular conditions, SUMOylation of IE2 was severely affected in the phospho-deficient variant. Hence, phosphorylation by CK2 increases both non-covalent and covalent interaction between IE2 and SUMO.

The covalent interaction between IE2 and SUMO is vital for IE2’s auto-repression activity (Reuter *et al*., 2018). SUMOylated IE2 can recruit chromatin modifiers, e.g., HDAC and HMTs, to repress its promoter. Phosphorylation by CK2 enhances IE2 SUMOylation and consequently, facilitates auto-repression. The non-covalent interaction with SUMO is essential to recruit SUMOylated transcription factors (e.g., TAF12) during transactivation (Kim *et al*., 2010). The functional significance of CK2-mediated phosphorylation of IE2-SIM1 is highlighted by the fact that the transactivation activity of phospho-deficient mutant pmIE2 decreased by 25% in comparison to wt-IE2. While phosphorylation of the Serine-rich region depletes the transactivation activity of IE2, phosphorylation of IE2-SIM1 in fragment 5B enhances the transactivation activity, indicating that PTMs regulate the complex functional interplay of IE2 domains.

Intriguingly, the activity of phospho-deficient pmIE2 is comparable to SIM mutated smIE2, suggesting that the effect of phosphorylation is equivalent to the presence of SIM in cellular conditions. Since the IE2-SIM1/SUMO interaction is weak, the tighter binding upon phosphorylation may be critical to ‘switch on’ the interaction with SUMOylated transcription factors and enable efficient transactivation. Global SUMOylation, including the SUMOylation of transcription factors, increase during HCMV infection (Scherer *et al*., 2016). The surge in SUMOylated transcription factors and the phosphorylation of viral transactivator IE2 by tegument CK2 may work in tandem to regulate the transcription of the viral genome.

A comparison of K_m_ values obtained from the kinetic analysis indicated that the presence of SIM reduced K_m_ by 2.5-fold, and phosphorylation reduced the K_m_ by another 3.5-fold. Together, the presence of SIM and its phosphorylation decreased the K_m_ between the substrate and the enzyme by 8-fold, which drastically enhances the rate of the SUMOylation. Herein, we have uncovered that the HCMV transactivator protein IE2 hijacks a cross talk between two host PTMs; SUMOylation and phosphorylation. The intriguing mechanism of phosphorylation-enhanced SUMO interaction improves the transactivation and auto-repression activity of the viral protein. A better understanding of the molecular mechanisms underlying viral hijack of the cross-talk between multiple PTMs might provide novel opportunities for intervention.

## Acknowledgment

The NMR data were acquired at the NCBS-TIFR NMR Facility. The authors would like to thank Purushotham Reddy for helping with NMR data acquisition. R.D. is the recipient of Ramalingaswamy fellowship from Department of Biotechnology (DBT), Government of India. This research was funded by intramural grants from the National Center for Biological Sciences, Tata Institute of Fundamental Research.

## Material and Methods

### Plasmid and Peptides

HCMV IE2 peptides were synthesized from Lifetein. pET28- SUMO1(C-His), pET28-SUMO2(C-His), pET11-AOS1/UBA2 and pET28-ΔN364 SENP2 were gifted by Dr. Christopher Lima, Sloan Kettering Institute, New York. pET28-UBC9 was obtained from Addgene (25213). This construct was used as a template for site-directed mutagenesis to obtain H20D UBC9, K14R UBC9, and C93K UBC9. SUMO1-pQE80L and SUMO2- pET15b were obtained from TIFR, Mumbai. Mammalian expression vectors, pDEST-SG5 HA-IE2, pUL54-Luc, and pSG5 Flag-SUMO1, were kind gifts from Dr. Jin-Hyun Ahn, Sungkyunkwan University School of Medicine. The pDEST-SG5 HA-IE2 was used as a template for site-directed mutagenesis to generate CIVI199/200/201/202AAAA HA-IE2 and S 203/205 A HA-IE2.

### Protein purification

All the proteins were expressed and purified from BL21 (DE3) cells in ^15^NH_4_Cl-M9 medium for labeled proteins and in LB for unlabelled proteins. Labeled proteins were purified in phosphate buffer for NMR, while in Tris buffer for in-vitro assays. ^15^N labeled SUMO1/2 were cultured at 37°C to 0.8 OD_600_ and induced with 0.5 mM IPTG for 4-5 hour. Cells were lysed by sonication in lysis buffer containing 50mM Na2HPO4 pH 8.0, 20 mM imidazole and 300 mM NaCl. Lysate was clarified by centrifugation and supernatant was incubated with pre-equilibrated Ni-NTA beads for 1 hour. After incubation, flow through was collected and beads were washed with at least 5 column volume lysis buffer. Protein was eluted by increasing concentration of imidazole in lysis buffer. Fractions containing SUMO1/2 were concentrated and were further processed through a gel-filtration column (Superdex 75 16/600) in PBS buffer.

For in-vitro assays, E1 (UBA2/AOS1) and E2 (UBC9) were purified with Ni NTA affinity purification followed by Gel filtration as discussed above, but in Tris buffer. Lysis/wash buffer composition was 50 mM Tris pH 8, 350 mM NaCl, 1 mM PMSF, 1 mM beta-mercaptoethanol, 20 mM imidazole. Elution buffer contained 25 mM Tris pH8, 150 mM NaCl, 1 mM beta-mercaptoethanol, 250 mM imidazole and gel filtration buffer contained 20 mM Tris pH8, 50 mM NaCl, 1 mM beta-mercaptoethanol.

For SUMOylation assays, the mature form of SUMO1 and SUMO2 were obtained by processing CHis-SUMO1 or CHis-SUMO2 with SENP2. After Ni NTA purification, fractions containing CHis-SUMO1/2 were pooled up and incubated with purified His-ΔN 364 SENP2 (1:1000 molar ratio) at room temperature till complete digestion of C terminal extension. SENP2 and unprocessed SUMO were removed by passing the reaction mixture through Ni NTA beads, and flow-through containing mature SUMO was collected, concentrated and further purified with Superdex 75. ΔN 364 SENP2-pET 28b was expressed in BL21 (DE3) and purified through Ni NTA affinity purification as mentioned above. The buffer used for SENP2 purification is similar to the buffer used for E1 purification. After Ni-NTA, partially purified SENP2 was concentrated and stored.

### In-vitro biochemical assays

For SUMOylation of IE2-NTD or IE2-ppNTD or IE2-NTDm, 5 μM peptide and 5 μM SUMO1/2 were incubated with 1 μM E1 and 2.5 μM E2. The reaction was started by adding ATP. SUMOylation buffer contains 20 mM HEPES pH 7.5, 50 mM NaCl, 5 mM MgCl_2_, 0.1% Tween 20. The reaction was analyzed either on 12% SDS PAGE or by a change in anisotropy. Gels were imaged for FITC fluorophore (λ_ex_- 495 nm, λ_em_ - 519 nm). While anisotropy was measured using MOS450 fluorimeter (λ_ex_- 470 nm, λ_em_- 520- 560nm).

IE2-ppNTD was obtained by phosphorylating IE2-NTD with purified CK2. Human CK2 was obtained from NEB (P6010S). IE2-NTD was phosphorylated in buffer provided by the manufacturer which contains 50mM Tris pH 7.5, 10mM MgCl_2_, 0.1mM EDTA, 2mM DTT and 0.1% Brij35. In 50 μl reaction, 50 unit of CK2 was sufficient to phosphorylate 50 μM IE2 peptide in 2 hours at 30 °C. CK2 was heat inactivated at 65 °Cfor 15 min after IE2 phosphorylation. IE2-NTD phosphorylation was analysed using autoradiography or by MALDI-TOF mass spectrometry. Phosphorylated IE2-NTD (IE2-ppNTD) was used as a substrate for SUMOylation assays.

C93K UBC9-SUMO1 conjugates were made either with ^15^N-C93K UBC9 or ^15^N-SUMO1. Conjugation reaction was performed in buffer containing 20 mM CAPS (pH9.5), 50 mM NaCl, 5 mM MgCl_2_, 3 mM DTT in presence of 10 μM E1, 300 μM C93K UBC9, 500 μM SUMO1 and 3 mM ATP. The reaction was performed overnight at 37 °C. Conjugate formation was analysed on a SDS gel before further purification by MonoQ and SEC. Conjugation reaction mixture was diluted by 25 mM Tris (pH8) to reduce salt concentration. The diluted reaction mixture was loaded onto the MonoQ column and proteins were eluted with increasing concentration of NaCl (up to 1 M). Fractions containing C93K UBC9-SUMO1 conjugate were pooled and further purified by SD75 in PBS buffer.

Single turn over reactions was performed for Michaelis–Menten kinetics experiments. UBC9∼K11R SUMO2 conjugates were formed in a 50 μl reaction with 1 μM E1, 10 μM E2 and 10 μM K11R SUMO2 in SUMOylation buffer. The reaction was started by adding E1and was incubated at 37°C for 10min. The reaction was stopped by adding 1450 μl quenching buffer. Quenching buffer contained 20 mM HEPES pH 7.5, 50 mM NaCl, 5 mM EDTA, 0.1% Tween 20. IE2-NTD SUMOylation was performed by adding an equal volume of quenched reaction to different concentration of serially diluted IE2-NTD peptides (so that conjugates and peptides are further diluted by half to give desired concentration). Aliquots were taken at desired time points and the reaction was stopped by SDS loading dye. The reaction was resolved on 12% SDS PAGE in non-reducing conditions which was then transferred onto PVDF membrane and was blotted with SUMO2 antibody.

### NMR experiments

The NMR spectra of SUMO1 and SUMO2 were recorded at 298K on 800 MHz Bruker Avance III HD spectrometer with a cryoprobe head, processed with NMRpipe (Delaglio *et al*., 1995) and analyzed with Sparky(Kneller and Kuntz, 1993). All the NMR experiments with FHA-Chk2 were performed at 293K. The SUMO1/2 samples were prepared in PBS buffer, with 5 mM DTT at pH 7.4 and 10% D_2_O. ^1^H-^1^H TOCSY and ^1^H-^1^H NOESY were acquired and used to assign IE2-ppSIM. For NMR titration experiments, ∼3 mM peptides were titrated into ∼0.3 mM ^15^N-SUMO1 or ^15^N-SUMO2. The titration data was fit in 1:1 protein:ligand model using the equation CSP_obs_ = CSP_max_ {([P]_t_+[L]_t_+K_d_) - [([P]_t_+[L]_t_+K_d_)^2^- 4[P]_t_[L]_t_]^1/2^}/2[P]_t_, where [P]_t_ and [L]_t_ are total concentrations of protein and ligand at any titration point. The SUMO/IE2-ppSIM1 NMR samples were prepared in PBS buffer, with 5 mM DTT at pH 7.4 and 10% D_2_O. ^13^C, ^15^N half-filtered NOESY was collected on a ^13^C, ^15^N-SUMO1/IE2-ppSIM1 (1:1.5) complex sample with a mixing time of 200ms to measure intermolecular NOEs between SUMO1 and IE2-ppSIM1. A similar experiment was performed to obtain intermolecular NOEs in the SUMO2/IE2-ppSIM1 complex.

### Structure Determination and Modelling

Unambiguous restraints between the SUMO1 and IE2-ppSIM1 were determined from the intermolecular observed NOEs. Dihedral angles of IE2-ppSIM1 determined from 1H-1H TOCSY experiments were used to determine a structure of IE2-ppSIM1. The solution structure was calculated in HADDOCK (Dominguez, Boelens and Bonvin, 2003) using the structure of SUMO1 (PDB ID: 4WJO) and the extended structure of IE2-ppSIM1. Rigid body energy minimization generated one thousand initial complex structures, and the best 200 by total energy were selected for torsion angle dynamics and subsequent Cartesian dynamics in an explicit water solvent. Default scaling for energy terms was applied. The interface of SUMO1 was kept semi-flexible during simulated annealing and the water refinement steps. Following the standard benchmarked protocol, cluster analysis of the 200 water-refined structures yielded a single clear ensemble. The SUMO2/IE2-ppSIM1 complex was similarly docked, except that intermolecular restraints between SUMO2 and IE2-ppSIM1 were used and the starting structure of SUMO2 was taken from a crystal structure of free SUMO2 (PDB ID: 1WM3). The structures of IE2-SIM1/SUMO1 and IE2-SIM1/SUMO2 are deposited in the PDB with accession codes xxx and xxx.

The model of UBC9∼SUMO1/IE2-NTD was calculated in xplor-NIH. The restraints of UBC9∼SUMO1 were obtained from the PDB: 1Z5S. IE2-NTD was flexible. The other restraints used were the SUMO1/IE2-ppSIM1 noe restraints and UBC9/IE2-NTD CSPs. The distance of acceptor lysine K180 and the active site was restrained to 1.8Å. A hundred structures were calculated, and the ten lowest energy structures were used for analysis. The model of SUMO1/UBC9∼SUMO1/IE2-NTD, where an additional SUMO1 molecule is non-covalently bound to UBC9 was calculated similarly, except the SUMO1/UBC9 intermolecular restraints were estimated from the PDB: 2YZZ and the SUMO1/IE2-SIM1 restraints were set between the non-covalent bound SUMO1 and IE2-SIM1. The model of SUMO1- UBC9∼SUMO1/IE2-NTD complex, where the SUMO1 is covalently bound to UBC9, was calculated similarly, except that SUMO1-UBC9 restraints were obtained from PDB: 2VRR and the SUMO1/IE2-SIM1 restraints were set between the covalent bound SUMO1 and IE2- SIM1.

### Cell culture and transfection

HEK293T cells were maintained in DMEM with 10% serum. For any experiment, cells were seeded into 12 well tissue culture plates. Cells were transfected at 60-80% confluency with 500 ng Flag-SUMO1 and 500 ng HA-IE2 (wt or mutant as mentioned) using Lipofectamine 3000 reagent. Cells were harvested 48-hours post-transfection and lysed with 2x SDS loading dye. Lysates were run on 12% SDS gel and were probed with HA antibody (CST-3724) after blotting.

For Transactivation assays, HEK293Tcells were seeded into a 12 well plate and were cultured to 70-80% confluency. Cells were transfected with 100ng pUL54-Luc, 5 ng pTK-renilla (transfection control) and 400 ng wt-IE2 with or without 400ng Flag-SUMO1. In the case of mutants (Ser203, 205A-IE2 and 199CIVI202, AAAA-IE2) transfected plasmid amount was increased to acquire similar expression as wt-IE2. Luciferase concentration was measured 36-40 hour post-transfection by Dual-Glo-luciferase kit (Promega).

HA-IE2 was immuno-precipitated form HEK293T for PTM analysis by Mass spectrometer. One 100mm dish was transfected with 10µg HA-IE2 plasmid with Lipofectamine 3000. IP was performed 36-hour post-transfection. HA-tag Sepharose beads (CST) were used for pull down. The protocol provided by the manufacturer was followed for the IP. After IP, beads were directly loaded onto reducing SDS PAGE, and immunoprecipitated proteins were resolved. The band, matching the size of the protein of interest, was excised and analyzed by Mass Spectrometry.

## Supplementary Information Legends

**Figure S1.** (A) CSPs of NMR titration between 15N-SUMO1/IE2-SIM3. (B) Overlay of the ^15^N-edited HSQC spectra of free ^15^N-SUMO2 (red) with different stoichiometric ratios of IE2- SIM1 as given in the top left-hand side of the spectra. (C) Two regions of the spectra are expanded to show a shift of SUMO2 resonances during titration. (D) The CSPs in SUMO2 upon binding to IE2-SIM3.

**Figure S2.** The fit of SUMO1 peak shifts against the concentration ratio [IE2-SIM1]/[SUMO1] yielded the K_d_ of the SUMO1/IE2-SIM1 complex. The fit of nine typical residues is shown. The residues are labeled at the bottom right corner of each window.

**Figure S3.** The fit of SUMO2 peak shifts against the concentration ratio [IE2-SIM1]/[SUMO2] yielded the K_d_ of the SUMO2/IE2-SIM1 complex. The fit of nine typical residues is shown. The residues are labeled at the bottom right corner of each window.

**Figure S4.** Detected phosphorylation sites in IE2. The unphosphorylated residues are colored blue. The phosphorylated sites are colored either black, red or green. MAPK predicted sites are colored in green, CK2 predicted sites are colored in red and rest is colored in black. The SIM and SUMOylation sites are highlighted with a yellow background.

**Figure S5.** (A) CSPs of NMR titration between 15N-SUMO1/IE2-ppSIM1. (B) The fit of SUMO1 peak shifts against the concentration ratio [IE2-ppSIM1]/[SUMO1] yielded the K_d_ of the SUMO1/IE2-ppSIM1 complex. The fit of nine typical residues is shown. The residues are labeled In the bottom right corner of each window.

**Figure S6.** (A) CSPs of NMR titration between 15N-SUMO2/IE2-ppSIM1. (B) The fit of SUMO2 peak shifts against the concentration ratio [IE2-ppSIM1]/[SUMO2] yielded the K_d_ of the SUMO2/IE2-ppSIM1 complex. The fit of nine typical residues is shown. The residues are labeled In the bottom right corner of each window.

**Figure S7.** (A) The twenty lowest energy structure of SUMO1/IE2-ppSIM1 complex. (B) Selected strips from the ^13^C, ^15^N half-filtered NOESY spectra depicting intermolecular NOEs between ^13^C-bonded protons of ^13^C, ^15^N-labeled SUMO2, and unlabeled IE2-ppSIM1. ^13^C and ^1^H assignment of SUMO2 atoms are given on the right and left of the strips, respectively. The protons of IE2-ppSIM1 that show NOEs to SUMO2 are assigned. (C) The twenty lowest energy structure of SUMO2/IE2-ppSIM1 complex.

**Figure S8.** (A) The products of the SUMOylation reaction with SUMO2 and IE2-NTD as the substrate is resolved on the SDS-PAGE gel and imaged with a filter at 519 nm corresponding to FITC fluorescence. Bands of free IE2-NTD or conjugated with one, two or multiple (n) SUMO2 are marked. The time-points are given on the top of the gel. (B) Same as in (A) where IE2-NTDm replaced IE2-NTD. (C) SUMOylation of IE2-NTD monitored in real-time by Fluorescence anisotropy measurements. The –ATP experiment is a negative control, where IE2-NTD is not SUMOylated, and the fluorescence anisotropy does not change.

**Figure S9.** A representative graph of the detected IE2-NTD∼SUMO conjugates against time with various concentrations of IE2-NTD.

**Figure S10.** A) CSPs observed in ^15^N-UBC9(C93K) upon titration with IE2-NTD. The CSPs in the loop between the active site and α2, and in the loop between α2 and α3, that were observed in a similar titration in wt-UBC9, are absent here. B) CSPs observed in SUMO1 non-covalently bound to UBC9 and titrated with IE2-NTD. The major CSPs are between β2α1 region, which is the same as apo SUMO1/IE2-SIM1 interactions. The residues marked with asterisk disappear during titration.

**Figure S11.** (A) The ten lowest energy model structures of the UBC9∼SUMO1/IE2-NTD complex. (B) Same as in (A) for the SUMO1/UBC9∼SUMO1/IE2-NTD, where SUMO1 is non-covalently bound to UBC9. (C) Same as in for the SUMO1-UBC9∼SUMO1/IE2-NTD complex, where SUMO1-UBC9 denotes the SUMO1 covalently linked to K14 of UBC9. UBC9 is colored orange, thioester conjugated SUMO1 is colored purple, covalently/non-covalently bound SUMO1 is in pink, and IE2-NTD is colored cyan. IE2-SIM1 is colored red. K180 in IE2-NTD is colored blue. C-terminal Glycine 97 is conjugated SUMO1 is colored yellow. Active site cysteine in colored magenta.

**Figure S12.** (A) Model and lowest energy structure of the SUMO1/UBC9∼SUMO1/IE2-NTD complex, where SUMO1 is non-covalently bound to UBC9. Surface representations of UBC9, conjugated SUMO1, and non-covalently bound SUMO1 is shown. UBC9 is colored orange, conjugated SUMO1 is purple, non-covalently bound SUMO1 in pink, and IE2-NTD is colored in cyan. (B) Zoomed view of the active site in (A), where E178 forms a salt-bridge with K101. (C) Same as in (A) for the SUMO1-UBC9∼SUMO1/IE2-NTD complex, where SUMO1-UBC9 denotes the SUMO1 covalently linked to K14 of UBC9. (D) Zoomed view of the active site in (A), where E178 forms a salt-bridge with K74. IE2-SIM1 is colored red in all figures.

**Figure S13.** (A) The IE2-NTD∼SUMO and IE2-ppNTD conjugates were resolved on an SDS page gel and blotted with the anti-SUMO antibody with various concentrations of IE2-NTD. (B) The modified IE2-NTD∼SUMO were quantified against time with various concentrations of IE2-NTD.

**Figure S14.** Expression profile of wt IE2 and mutant IE2s used in the Luciferase transactivation assay.

**Figure S15.** Figure 5D is replotted here to show that IE2∼SUMO conjugates increase upon overexpression of SUMO. IE2∼SUMO has two bands. The lower band IE2∼SUMO is where IE2 is conjugated with endogenous SUMO. The upper band is where IE2 is conjugated with transfected FLAG3x-SUMO1.

