## Supplementary for "Phosphorylation by CK2 increases the SUMO-dependent activity of Cytomegalovirus transactivator IE2"

**[1]National Center for Biological Sciences, TIFR, Bangalore, India**

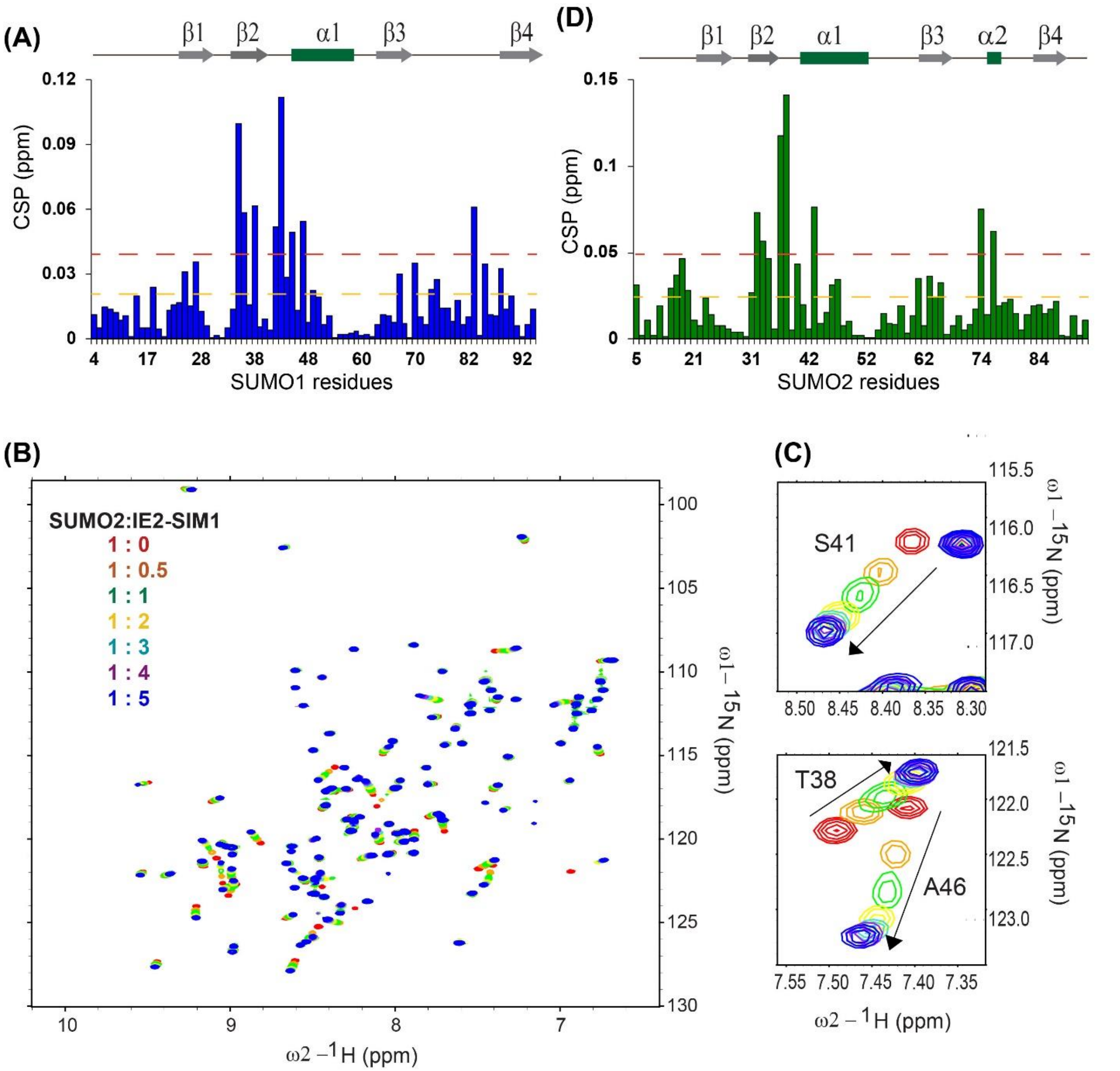


**Figure S1.** (A) CSPs of NMR titration between 15N-SUMO1/IE2-SIM3. (B) Overlay of the $^{15}$N-edited HSQC spectra of free $^{15}$N-SUMO2 (red) with different stoichiometric ratios of IE2-SIM1 as given in the top left-hand side of the spectra. (C)Two regions of the spectra are expanded to show a shift of SUMO2 resonances during titration. (D) The CSPs in SUMO2 upon binding to IE2-SIM3.

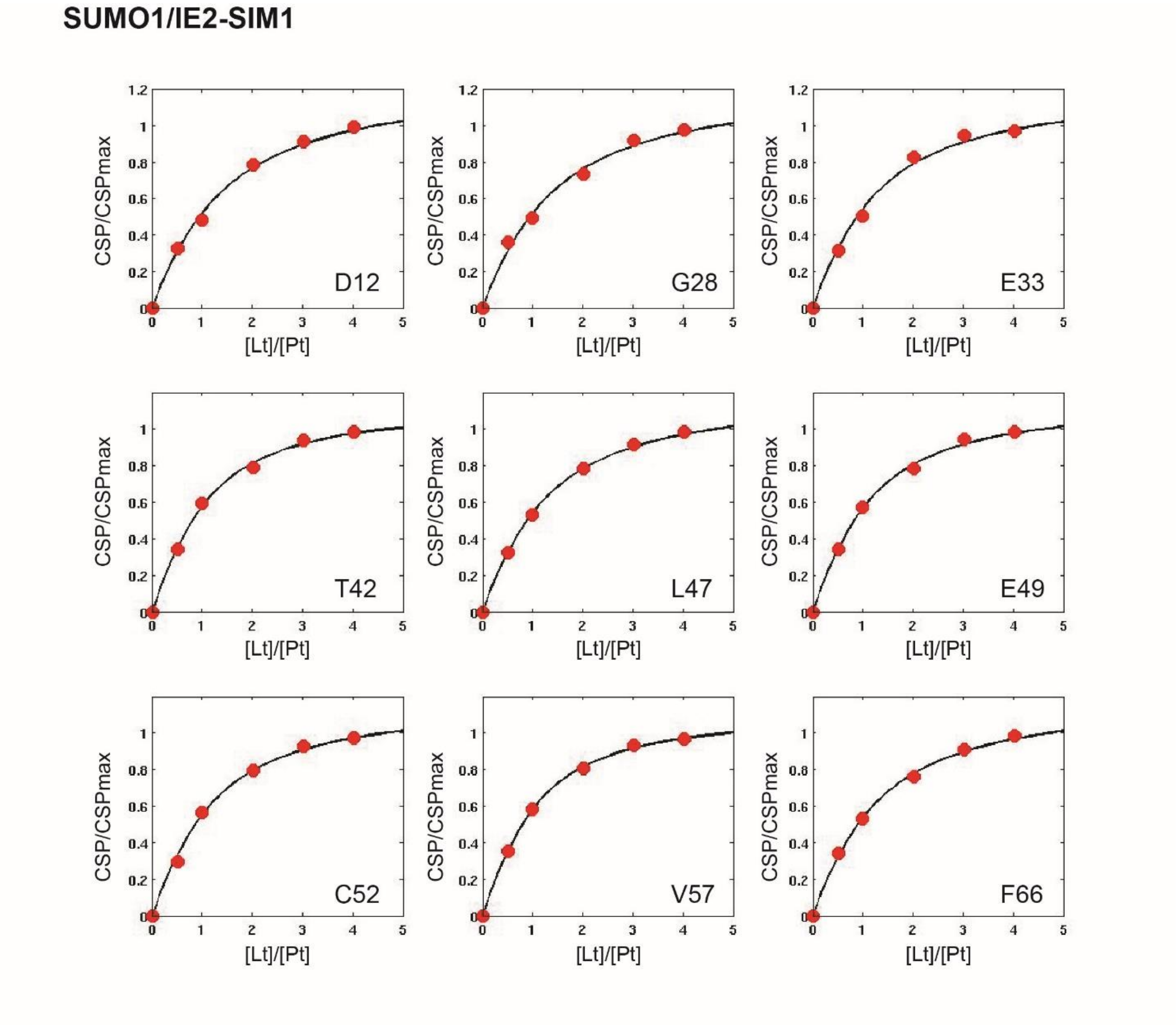


**Figure S2.** The fit of SUMO1 peak shifts against the concentration ratio [IE2-SIM1]/[SUMO1] yielded the $K_d$ of the SUMO1/IE2-SIM1 complex. The fit of nine typical residues is shown. The residues are labeled at the bottom right corner of each window.

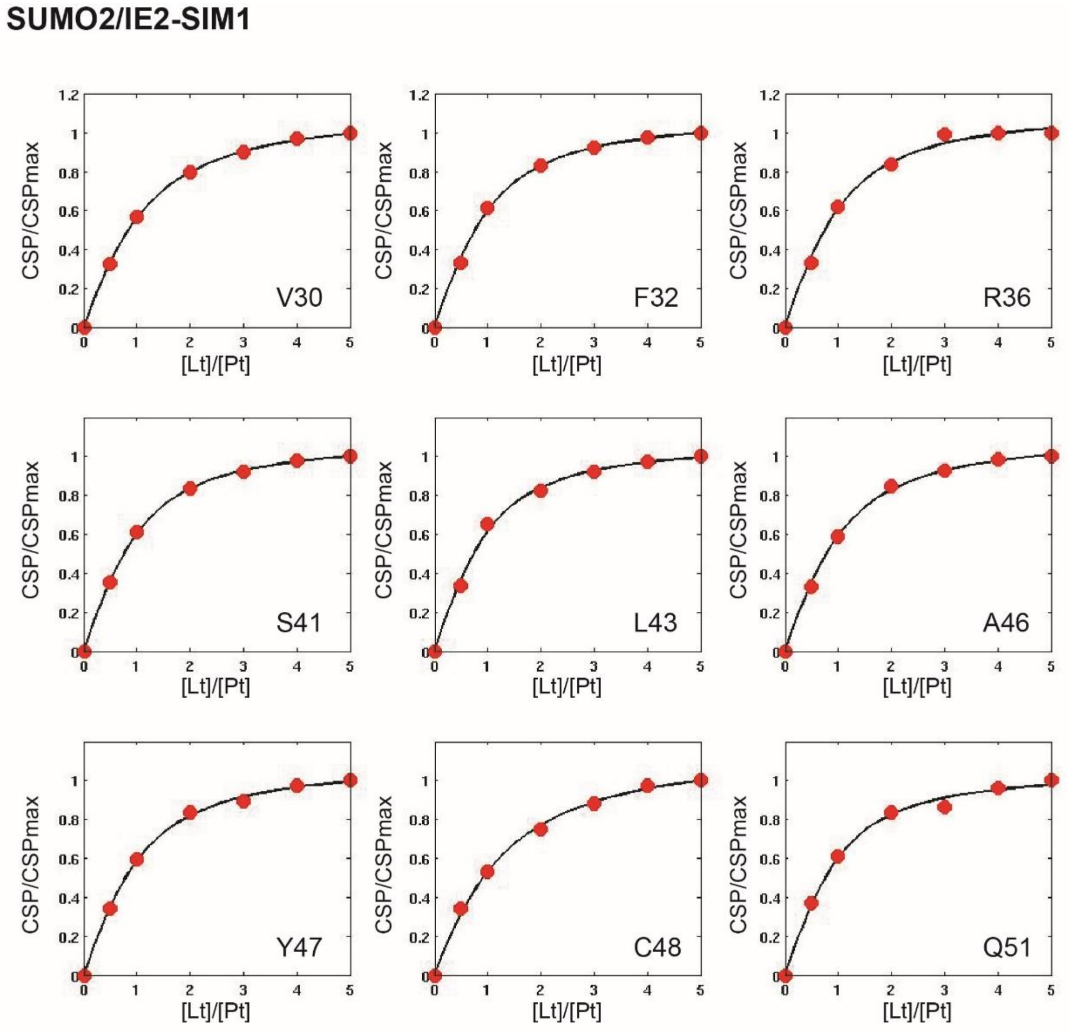


**Figure S3.** The fit of SUMO2 peak shifts against the concentration ratio [IE2-SIM1]/[SUMO2] yielded the $K_d$ of the SUMO2/IE2-SIM1 complex. The fit of nine typical residues is shown. The residues are labeled at the bottom right corner of each window.

MESSAKRKMDPDNPDEGPSSKVPRPETPVTKATTFLQTMLRKEVNSQLSLGDPLFPELAE 60

ESLKTFEQVTEDCNENPEKDVLAELGDILAQAVNHAGIDSSSTGPTLTTHSCSVSSAPLN 120

KPTPTSVAVTNTPLPGASATPELSPRKKPRKTTRPFKVIIKPPVPPAPIMLPLIKQEDIK 180

PEPDFTIQYRNKIIDTAGCIVISDSEEEQGEEVETRGATASSPSTGSGTPRVTSPTHPLS 240

QMNHPPLPDPLGRPDEDSSSSSSSSCSSASDSESESEEMKCSSGGGASVTSSHHGRGGFG 300

GAASSSLLSCGHQSSGGASTGPRKKKSKRISELDNEKVRNIMKDKNTPFCTPNVQTRRGR 360

VKIDEVSRMFRNTNRSLEYKNLPFTIPSMHQVLDEAIKACKTMQVNNKGIQIIYTRNHEV 420

KSEVDAVRCRLGTMCNLALSTPFLMEHTMPVTHPPEVAQRTADACNEGVKAAWSLKELHT 480

HQLCPRSSDYRNMIIHAATPVDLLGALNLCLPLMQKFPKQVMVRIFSTNQGGFMLPIYET 540

AAKAYAVGQFEQPTETPPEDLDTLSLAIEAAIQDLRNKSQ

**Figure S4.** Detected phosphorylation sites in IE2. The unphosphorylated residues are colored blue. The phosphorylated sites are colored either black, red or green. MAPK predicted sites are colored in green, CK2 predicted sites are colored in red and rest is colored in black. The SIM and SUMOylation sites are highlighted with a yellow background.

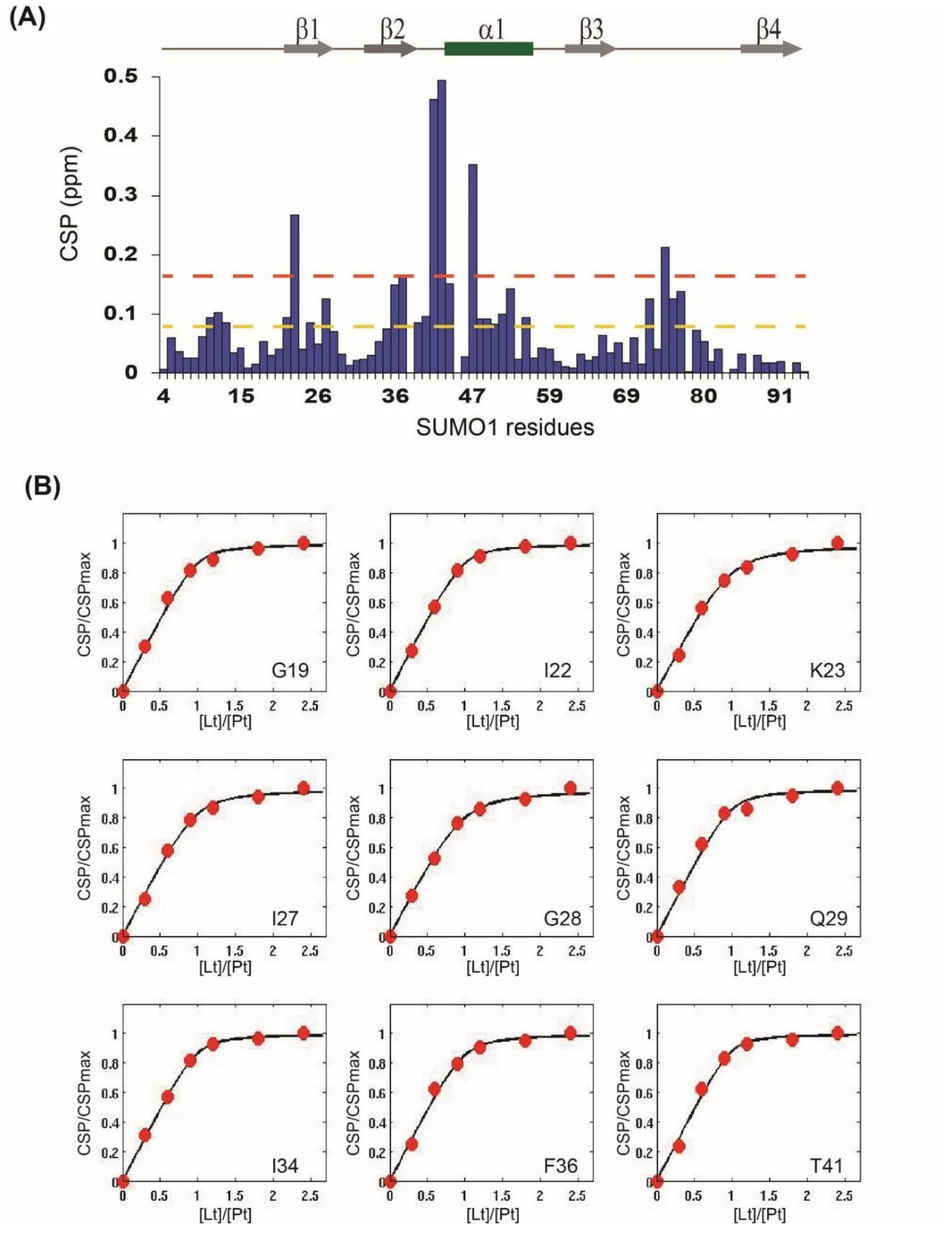


**Figure S5.** (A) CSPs of NMR titration between 15N-SUMO1/IE2-ppSIM1. (B) The fit of SUMO1 peak shifts against the concentration ratio [IE2-ppSIM1]/[SUMO1] yielded the $K_d$ of the SUMO1/IE2-ppSIM1 complex. The fit of nine typical residues is shown. The residues are labeled In the bottom right corner of each window.

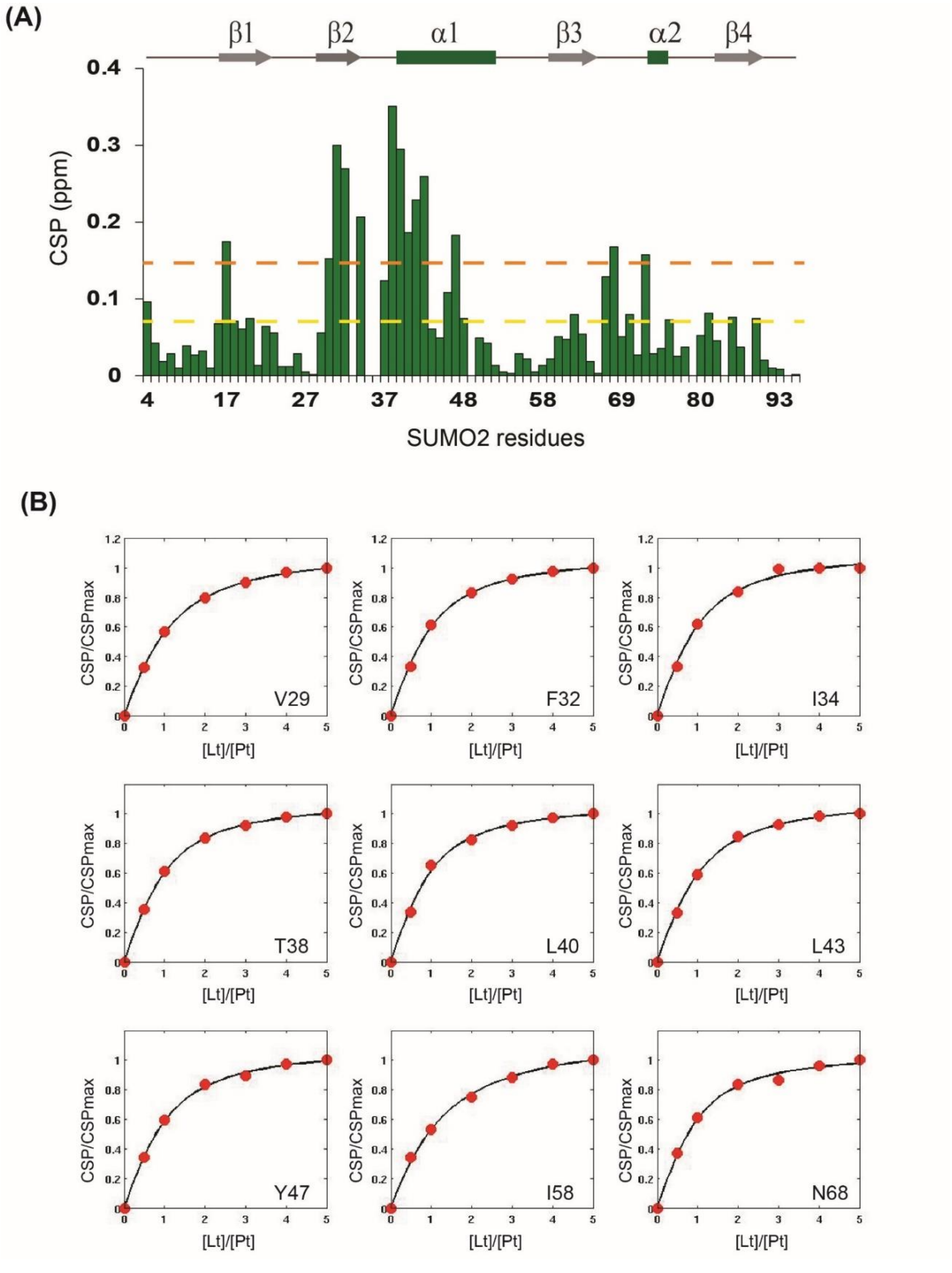


**Figure S6.** (A) CSPs of NMR titration between 15N-SUMO2/IE2-ppSIM1. (B) The fit of SUMO2 peak shifts against the concentration ratio [IE2-ppSIM1]/[SUMO2] yielded the $K_d$ of the SUMO2/IE2-ppSIM1 complex. The fit of nine typical residues is shown. The residues are labeled In the bottom right corner of each window.

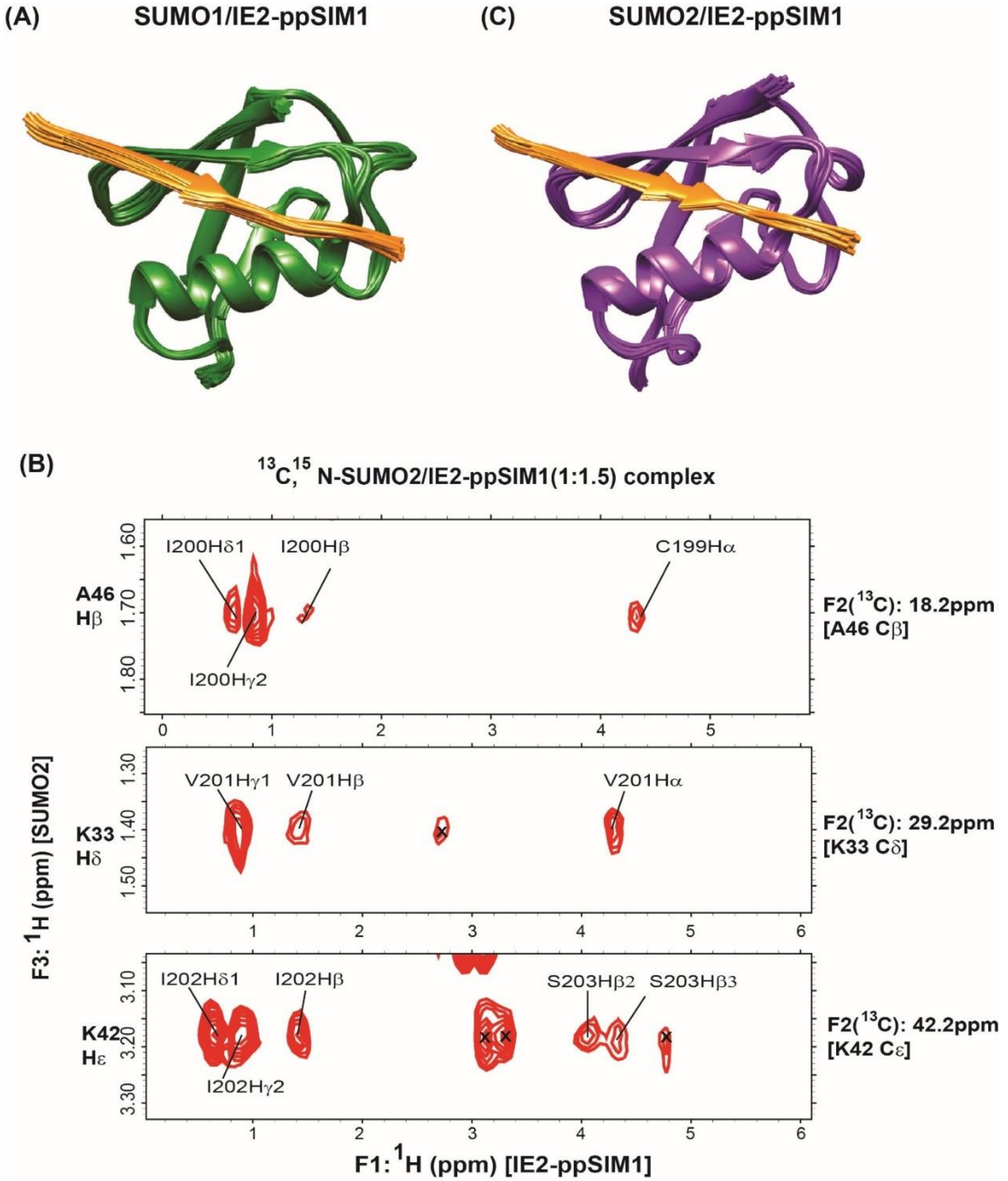


**Figure S7.** (A) The twenty lowest energy structure of SUMO1/IE2-ppSIM1 complex. (B) Selected strips from the $^{13}$C, $^{15}$N half-filtered NOESY spectra depicting intermolecular NOEs between $^{13}$C-bonded protons of $^{13}$C, $^{15}$N-labeled SUMO2, and unlabeled IE2-ppSIM1. $^{13}$C and $^{1}$H assignment of SUMO2 atoms are given on the right and left of the strips, respectively. The protons of IE2-ppSIM1 that show NOEs to SUMO2 are assigned. (C) The twenty lowest energy structure of SUMO2/IE2-ppSIM1 complex.

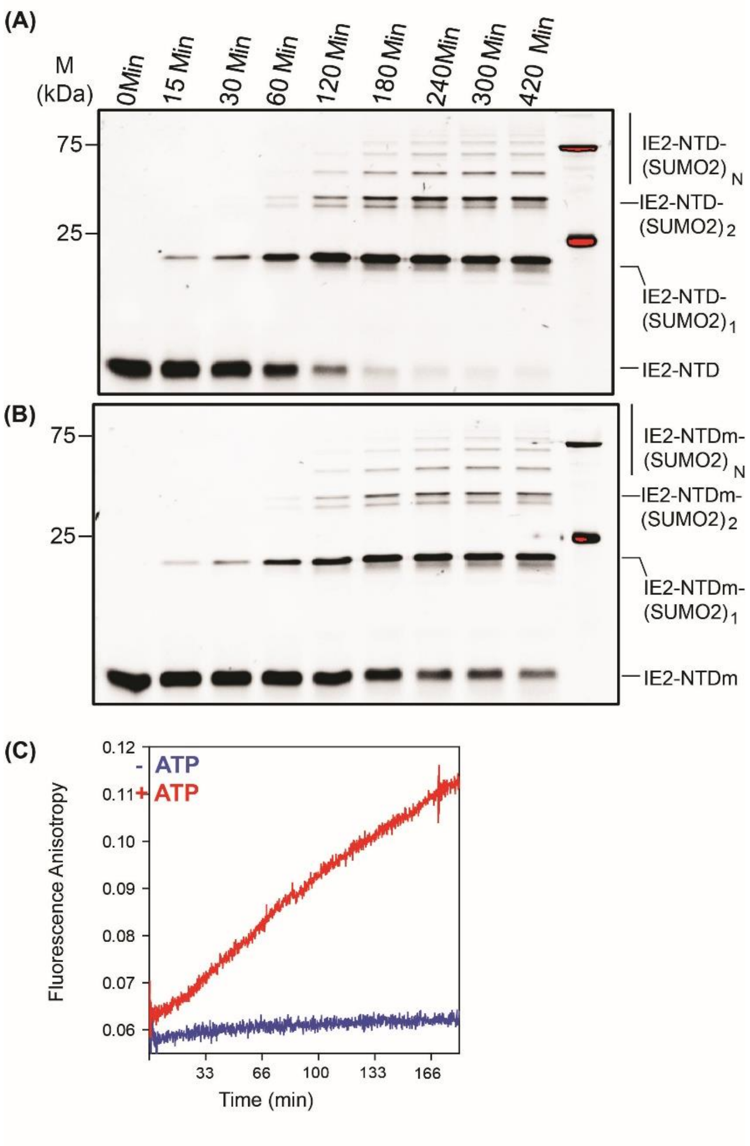


**Figure S8.** (A) The products of the SUMOylation reaction with SUMO2 and IE2-NTD as the substrate is resolved on the SDS-PAGE gel and imaged with a filter at 519 nm corresponding to FITC fluorescence. Bands of free IE2-NTD or conjugated with one, two or multiple (n) SUMO2 are marked. The time-points are given on the top of the gel. (B) Same as in (A) where IE2-NTDm replaced IE2-NTD. (C) SUMOylation of IE2-NTD monitored in real-time by Fluorescence anisotropy measurements. The –ATP experiment is a negative control, where IE2-NTD is not SUMOylated, and the fluorescence anisotropy does not change.

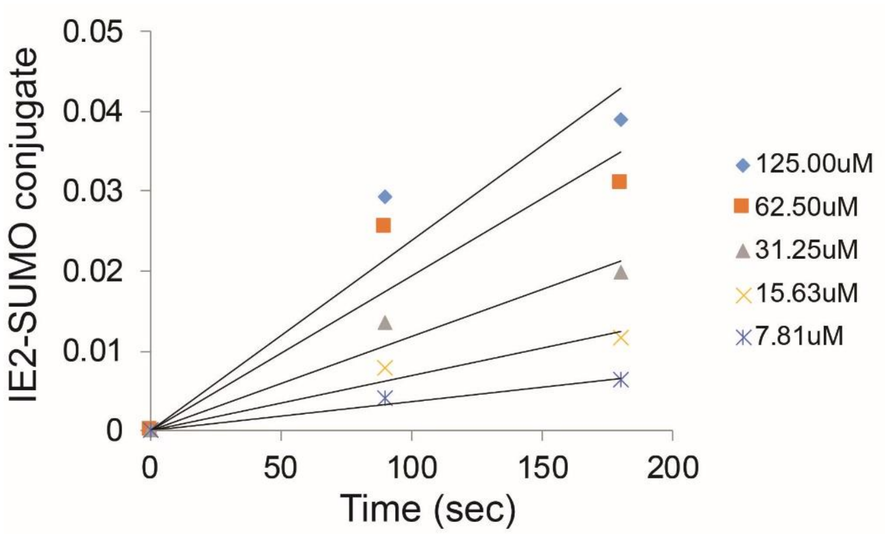


**Figure S9.** A representative graph of the detected IE2-NTD~SUMO conjugates against time with various concentrations of IE2-NTD.

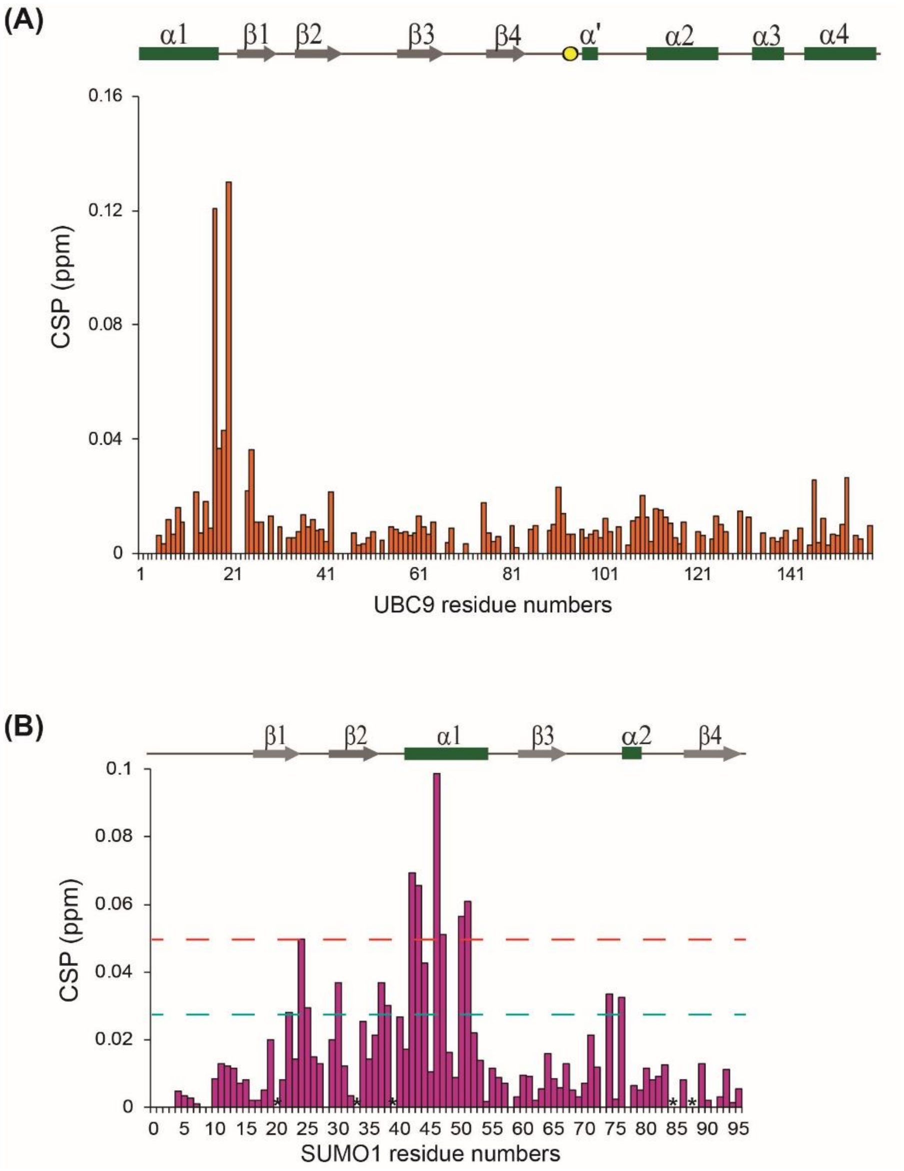


**Figure S10.** A) CSPs observed in $^{15}$N-UBC9(C93K) upon titration with IE2-NTD. The CSPs in the loop between the active site and α2, and in the loop between α2 and α3, that were observed in a similar titration in wt-UBC9, are absent here. B) CSPs observed in SUMO1 non-covalently bound to UBC9 and titrated with IE2-NTD. The major CSPs are between β2α1 region, which is the same as apo SUMO1/IE2-SIM1 interactions. The residues marked with asterisk disappear during titration.

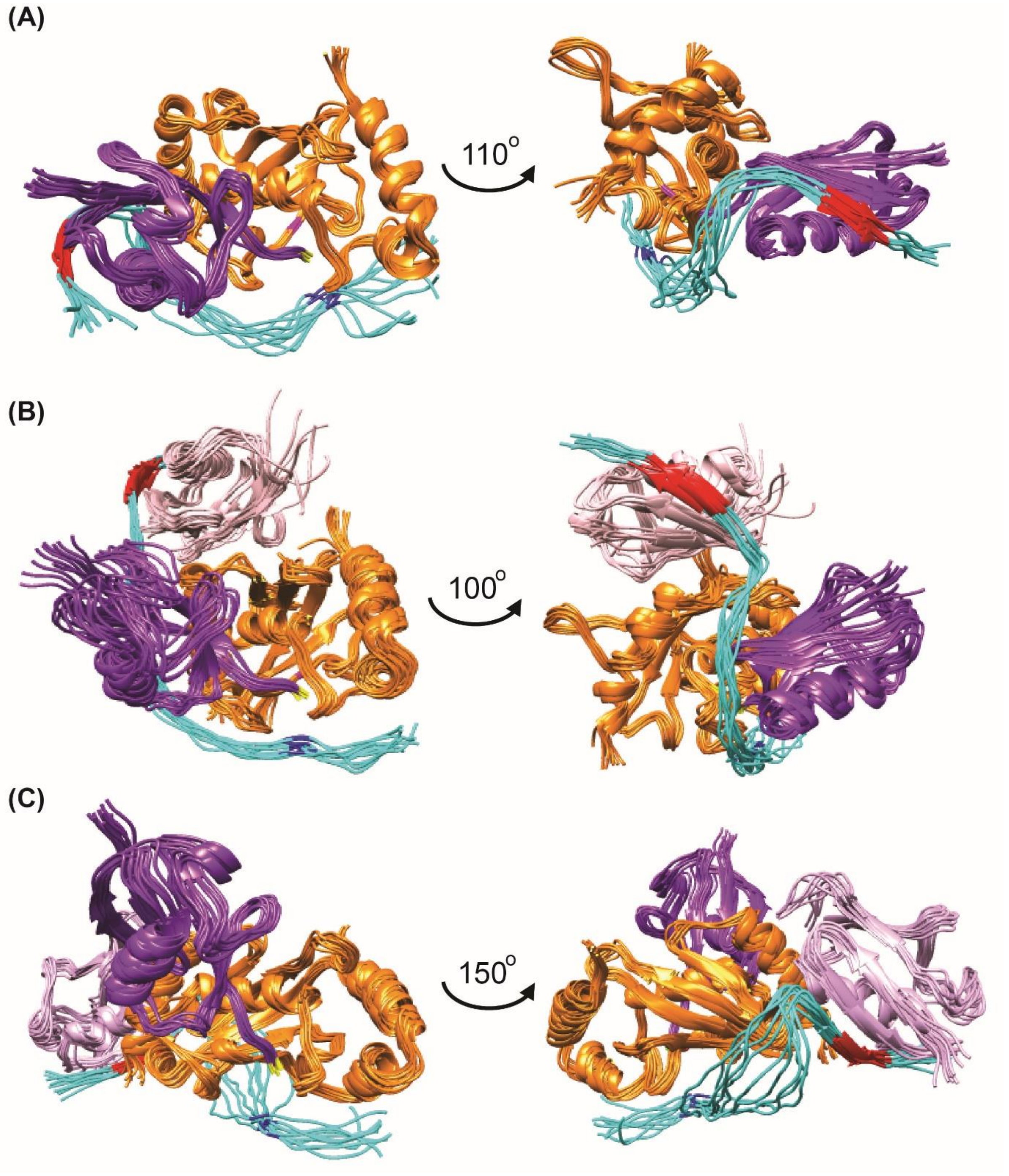


**Figure S11.** (A) The ten lowest energy model structures of the UBC9~SUMO1/IE2-NTD complex. (B) Same as in (A) for the SUMO1/UBC9~SUMO1/IE2-NTD, where SUMO1 is non-covalently bound to UBC9. (C) Same as in for the SUMO1-UBC9~SUMO1/IE2-NTD complex, where SUMO1-UBC9 denotes the SUMO1 covalently linked to K14 of UBC9. UBC9 is colored orange, thioester conjugated SUMO1 is colored purple, covalently/non-covalently bound SUMO1 is in pink, and IE2-NTD is colored cyan. IE2-SIM1 is colored red. K180 in IE2-NTD is colored blue. C-terminal Glycine 97 is conjugated SUMO1 is colored yellow. Active site cysteine in colored magenta.

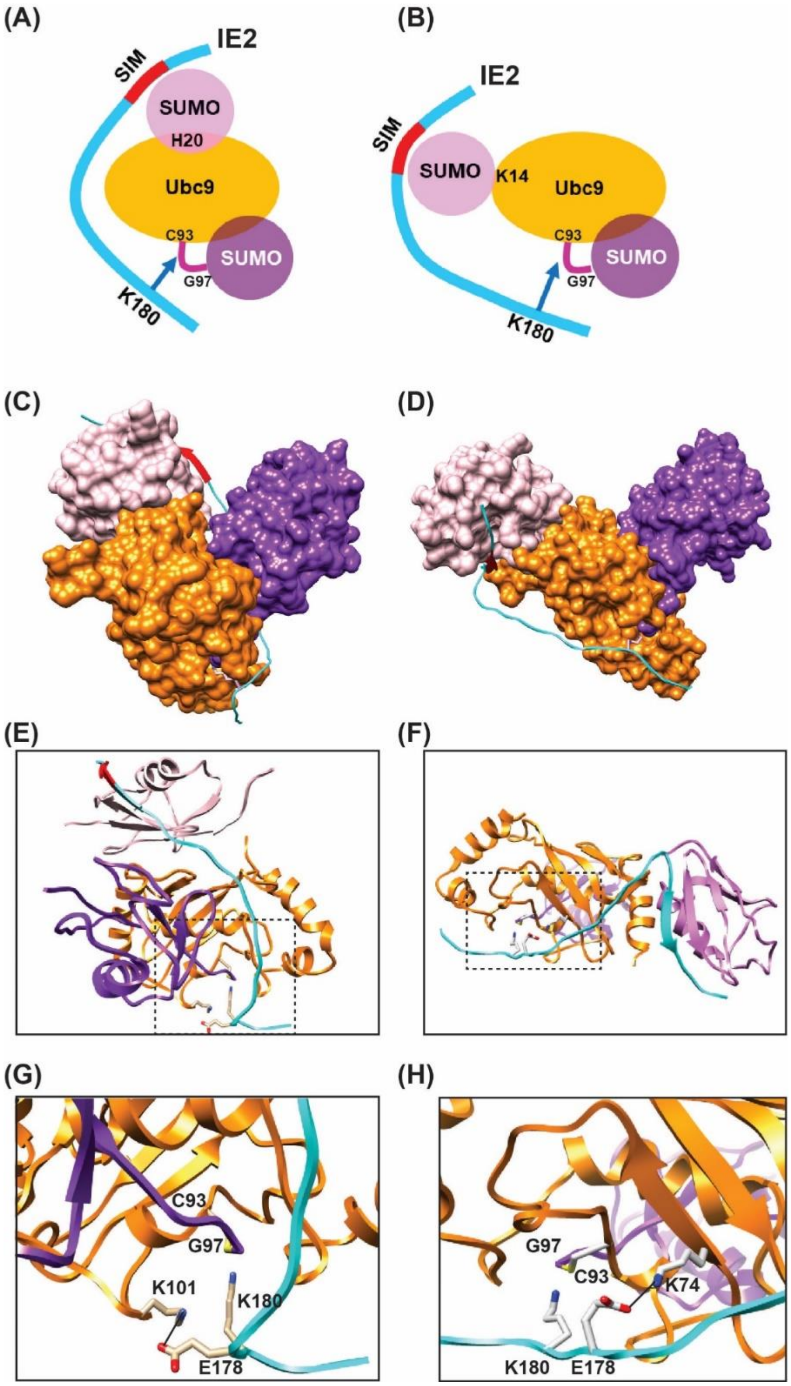


**Figure S12.** (A) Model and lowest energy structure of the SUMO1/UBC9~SUMO1/IE2-NTD complex, where SUMO1 is non-covalently bound to UBC9. Surface representations of UBC9, conjugated SUMO1, and non-covalently bound SUMO1 is shown. UBC9 is colored orange, conjugated SUMO1 is purple, non-covalently bound SUMO1 in pink, and IE2-NTD is colored in cyan. (B) Zoomed view of the active site in (A), where E178 forms a salt-bridge with K101. (C) Same as in (A) for the SUMO1-UBC9~SUMO1/IE2-NTD complex, where SUMO1-UBC9 denotes the SUMO1 covalently linked to K14 of UBC9. (D) Zoomed view of the active site in (A), where E178 forms a salt-bridge with K74. IE2-SIM1 is colored red in all figures.

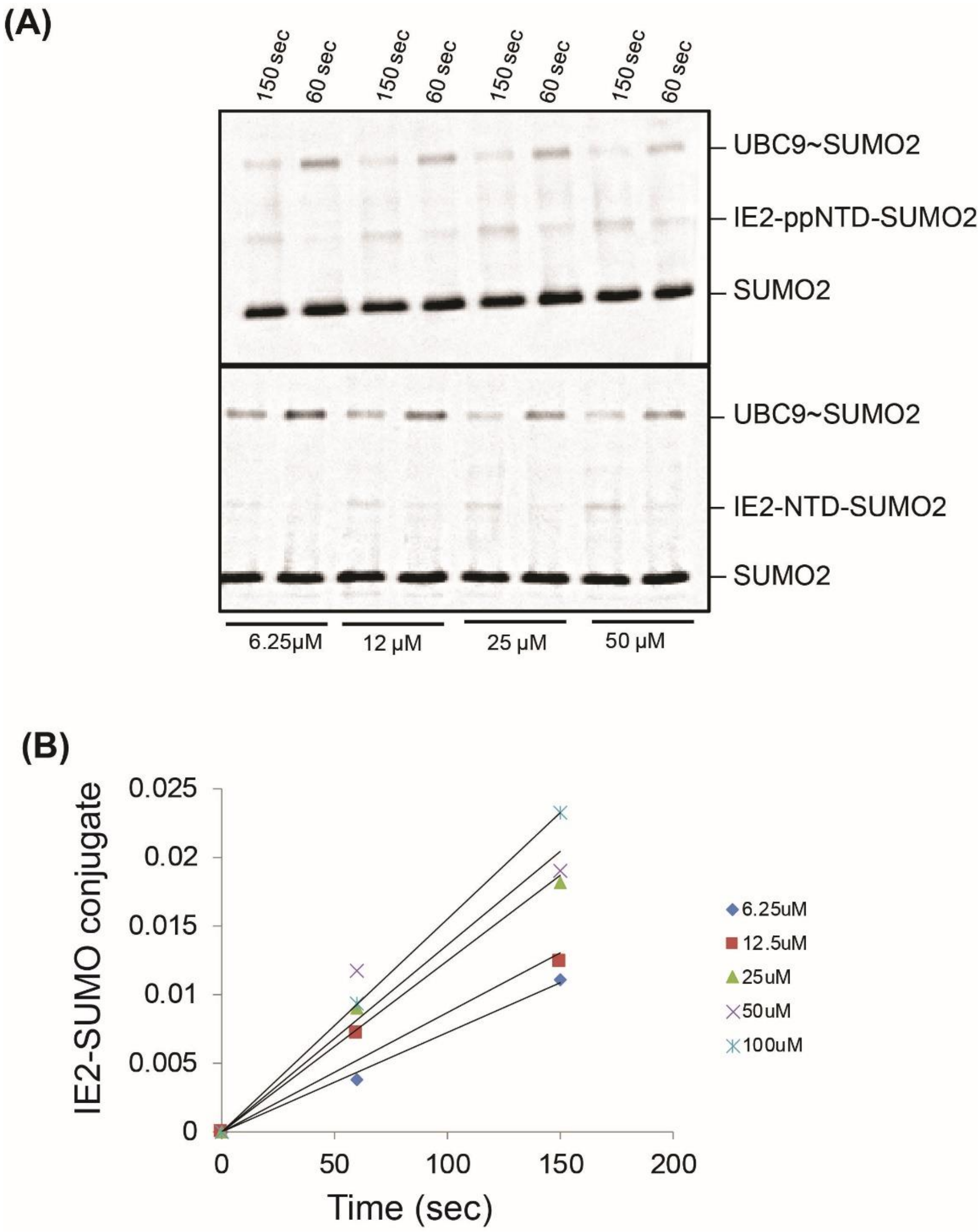


**Figure S13.** (A) The IE2-NTD~SUMO and IE2-ppNTD conjugates were resolved on an SDS page gel and blotted with the anti-SUMO antibody with various concentrations of IE2-NTD. (B) The modified IE2-NTD~SUMO were quantified against time with various concentrations of IE2-NTD.

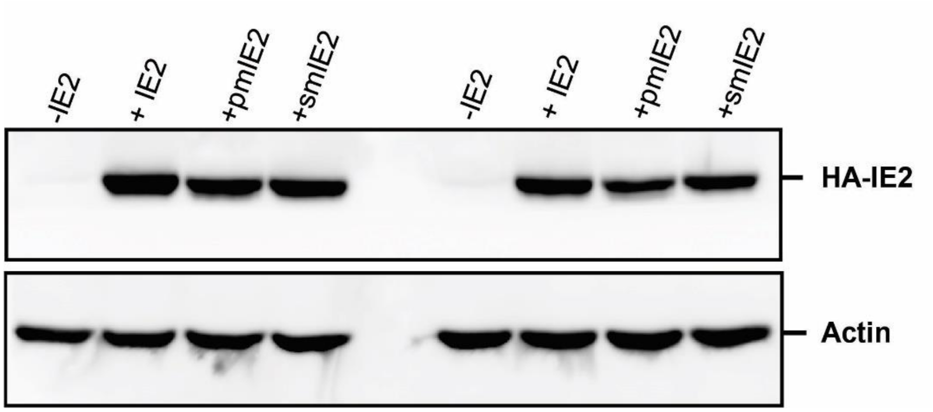


**Figure S14.** Expression profile of wt IE2 and mutant IE2s used in the Luciferase transactivation assay.

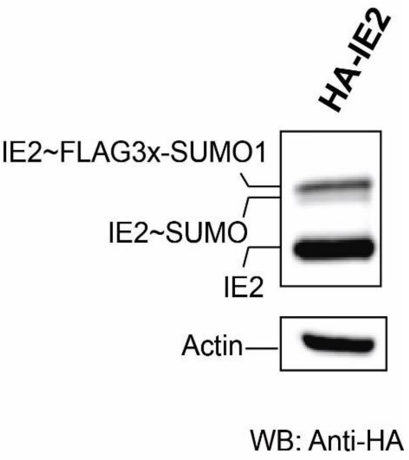


**Figure S15.** Figure 5D is replotted here to show that IE2~SUMO conjugates increase upon overexpression of SUMO. IE2~SUMO has two bands. The lower band IE2~SUMO is where IE2 is conjugated with endogenous SUMO. The upper band is where IE2 is conjugated with transfected FLAG3x-SUMO1.
